# Splicing of a core clock gene regulates seasonal adaptations by a winter gating mechanism

**DOI:** 10.1101/2025.11.06.687099

**Authors:** S. Hidalgo, R. Del Rio, C.A. Tabuloc, L.A. Perez Hernandez, A.L. Berry, Y.D. Cai, K.M. Lewald, J.C. Chiu

**Author notes:** Correspondence to: Sergio Hidalgo,; Joanna C. Chiu.

## Abstract

Seasons bring changes to the environment. Many organisms adjust their physiology and behavior in response to seasonal changes in order to survive. Although the molecular mechanisms mediating the integration of seasonal cues are still unclear, the working model indicates the involvement of the circadian clock. Notably, the circadian neuropeptide Pigment Dispersing Factor (PDF), an output of the circadian clock, has been shown to alter its expression and activity in response to seasonal changes to facilitate seasonal adaptations in insects. Here, we show that the alternative splicing (AS) of a circadian clock gene, *timeless (tim)*, regulates the seasonal responses of PDF signaling in *Drosophila melanogaster*. We first showed that *tim-sc,* the predominant isoform in winter, is regulated by photoperiod in addition to temperature, while the expression of the canonical *tim-l* isoform is primarily sensitive to temperature. We then demonstrated that *tim-sc* maintains physiology and behavior in a “winter lock” state by modulating PDF. At the cellular and molecular level, TIM-SC behavior differs from the canonical TIM-L. Interestingly, flies expressing *tim-sc* did not fully phenocopy wild-type flies reared in winter conditions, suggesting that other mechanisms are at play in regulating seasonal adaptations, despite the importance of *tim* AS.

## Introduction

Animals, plants, and even bacteria can anticipate and adapt to seasonal changes in environmental conditions^1–4^. These adaptations are diverse, involving a wide array of physiological, behavioral, and phenotypical changes such as developmental arrest (i.e., diapause)^5^, reproduction^3^, hibernation^6^, and migration^7–9^, among others. Seasonal changes allow organisms to endure long periods of extreme temperature conditions and limited food availability; thus, engaging in these changes needs to be precisely timed. Several phenological cues are used to inform seasonal environmental changes, with photoperiod (i.e., length of day) and temperature being the most prominent ones in temperate climates^10^. Little is known about how organisms integrate these cues to coordinate and establish seasonal programs in physiology and behavior, particularly in animals.

In the mid-1930s, Erwin Bünning proposed that the circadian clock, the mechanism that allows for maintaining daily rhythms in physiology and behavior, is indispensable for photoperiodism in plants^11,12^. Since then, studies have shown this to be the case in plants^13^ and in several vertebrate^14^ and invertebrate^15^ animals. In mammals and birds, the suprachiasmatic nucleus, the core circadian structure in the brain, integrates photoperiodic cues and regulates the rhythmic nocturnal release of melatonin from the pineal gland^1,16^. This initiates a hormonal cascade involving the thyroid-stimulating hormone (TSH) in the *pars tuberalis* of the pituitary gland and the regulation of tanycytes in the third ventricle of the hypothalamus, which triggers seasonal adaptations^17–24^. How circadian outputs, such as melatonin, are regulated in the seasonal context is still unclear, and exploring these questions in seasonal vertebrate models is still challenging given their genetic intractability. Thus, simple insect models have proven useful in this context.

In several insects, including *Drosophila melanogaster*, temperature has a massive influence on seasonal adaptations in addition to photoperiodic cues^25,26^. Both photoperiod and temperature are integrated by the circadian clock neuronal network (CCNN) to enable decisions on entering, maintaining, and exiting seasonal adaptations^27^. These neurons relay seasonal cues to neurosecretory cells in the *pars lateralis* (PL) and *pars intercerebralis* (PI), which in turn trigger seasonal adaptations by secreting hormone-like peptides^28^. The circadian neuropeptide Pigment Dispersing Factor (PDF) has been reported to play an important role in seasonality across insects^28–35^. PDF is a circadian output released by a group of circadian neurons within the CCNN named ventral lateral neurons (LNvs) and it coordinates the CCNN rhythmicity under constant darkness in *D. melanogaster*^36–39^. Under cold conditions, PDF levels drop, allowing the accumulation of the protein EYES ABSENT (EYA)^40^. PDF relays both photoperiod and temperature cues to cells in the PI to regulate seasonal adaptations, a feature that seems to be conserved in other insects, including mosquitoes, bugs, and flies ^32,41–45^. Since PDF is a circadian output, it is conceivable that the integration of seasonal cues occurs upstream in the circadian clock.

At the molecular level, the circadian clock functions as a transcriptional-translational feedback loop in which positive elements promote the transcription of negative elements that, once translated, block their own transcription in an exquisitely regulated process that takes around 24 hours^46^ . In *D. melanogaster*, the positive elements are *clock* (*clk*) and *cycle* (*cyc*), and the negative elements are *timeless* (*tim*) and *period* (*per*)^47–50^. Genetic manipulations by mutating or silencing these genes render a diverse array of phenotypes in several insects, highlighting the involvement of clock genes in seasonal adaptations (reviewed in ^29^). Nonetheless, how the circadian clock integrates seasonal cues is still unclear. In this regard, alternative splicing (AS) has arisen as a potential mechanism, given that multiple clock genes exhibit temperature-sensitive AS^51–53^.

AS is the process by which different isoforms are generated from the same pre-processed RNA. This mechanism mediates a wide array of regulations, including the generation of protein variants with altered structure and function, or controlling transcripts’ stability and degradation (reviewed in ^54^). AS is key to providing phenotypic plasticity and adaptation to the environment. As such, a role of AS in seasonal adaptations has been supported by evidence in several models, including plants and animals^55–59^. For instance, changes in the AS landscape have been observed in different tissues of hibernating brown bears^60^. In seasonal morphs of butterfly *Bicyclus anynana*, AS affects a subset of genes without changes in gene expression level^61^. In the Japanese quail, one particular isoform of *Eya3* containing exon 7 is expressed in the *pars tuberalis* under long days^62^, and AS affects the seasonal expression of the blue-light activated Cryptochrome in the retina of European Robins^63^.

Importantly, seasonal cues have been shown to modulate AS of core clock genes in several species. In *D. melanogaster*, AS of the 3’ terminal intron of *per* is observed under cold temperatures, in the lab and in natural conditions, which contributes to differential accumulation of *per* to regulate seasonal locomotor adaptations ^51,52,57^. Additionally, AS events have been shown to directly affect the structure and function of the core clock proteins. For instance, we recently described that *clk* undergoes temperature-dependent alternative splicing, which generates a CLK isoform with a 4 amino acid deletion adjacent to the CLK DNA binding domain, affecting CLK function in regulating circadian output and PDF response to temperature^64^. Another clock protein, TIM, undergoes even more pronounced AS-dependent changes in sequence/structure^65^ (Fig. 1A). Under cold conditions, two isoforms are generated, *tim-cold (tim-c)* and *tim-short-and*-*cold (tim-sc)*, encoding proteins differing in 33 and 507 amino acids, respectively, from the canonical TIM-long (TIM-L) isoform^66–69^. Expression of *tim-c* and *tim-sc* almost completely replaces *tim-l* in winter-like cold conditions, making these isoforms particularly interesting in the context of seasonal adaptations. Supporting this notion, *tim-*null mutants have impaired reproductive arrest response in response to seasonal changes^68^. Additionally, *tim* displays naturally occurring, clinally distributed alleles, *l* and *ls*, whose protein products exhibit differential light sensitivity^70–72^. All these data point to the potential role for *tim* in photoperiod and temperature cue integration.

**Figure 1.**
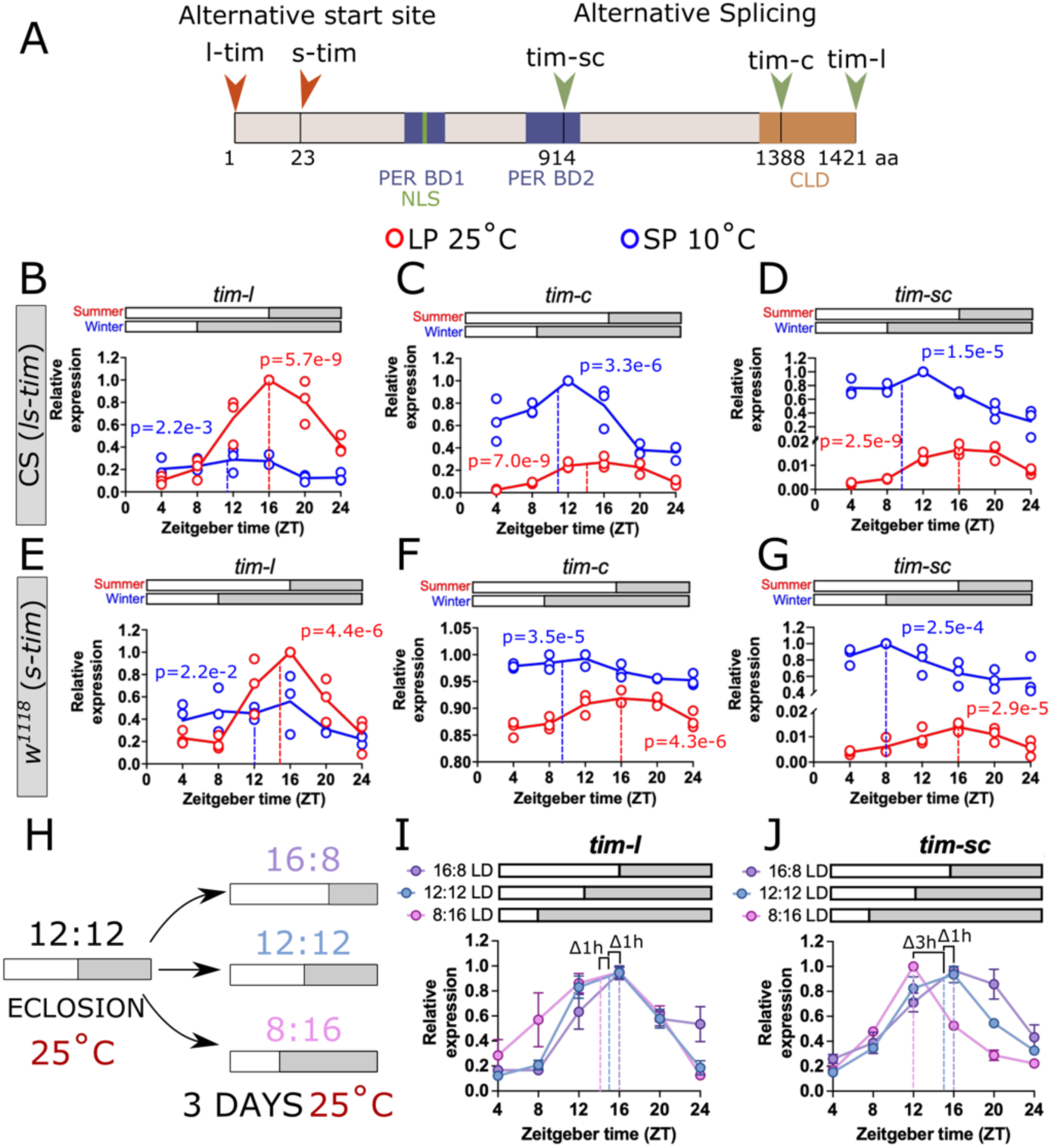
Messenger RNAs encoding temperature-dependent TIMELESS isoforms are differentially regulated by photoperiod. (A) Schematic of the TIMELESS protein. Alternative translation start sites from two naturally occurring alleles, *ls* and *s*, give rise to N-terminus variants, either L-TIM and S-TIM or just S-TIM, respectively. These two variants, L and S, vary in 23 amino acids. Temperature-dependent alternative splicing gives rise to two C-terminus major isoforms *tim-cold(c)* and *tim-shortcold(sc)*. Their protein products vary from the canonical TIM-Long(L) isoform in 33 and 507 amino acids, respectively. PER Binding domains 1 and 2 (PER BD1 and PER BD2), Nuclear localization Signal (NLS), and Cytoplasmic Localization Domain (CLD) are shown (reviewed in ^65^). Expression of (B) *tim-l*, (C) *tim-c*, and (D) *tim-sc* in summer-like, long photoperiod (LP; 16 hours light and 8 hrs dark cycles; 16:8 LD) at 25°C (red) and winter-like, short photoperiod (SP; 8 hours light and 16 hrs dark cycles; 8:16 LD) at 10°C (blue) in Canton-S (CS) flies, bearing the *ls* allele. Horizontal bars above the graphs depict light (white) and dark (gray) periods over the 24-hour cycle. Expression of (E) *tim-l*, (F) *tim-c*, and (G) *tim-sc* under the same conditions is shown for the *w*^1118^ flies bearing the *s* allele. (H) Experimental setting for changing photoperiod at constant temperature. Newly eclosed flies were moved to either LP (16:8 LD), equinox (12:12 LD), or SP (8:18 LD) at constant 25°C for 3 days before collection. Expression profiles of (I) *tim-l* and (J) *tim-sc* are shown for flies entrained under these conditions. A delta of 1 hr is observed across photoperiods in *tim-l,* while in *tim-sc* a 3 hr delta is observed in the peak expression between equinox and SP. Statistical p-values for rhythmicity obtained using CircaCompare are depicted in graphs B-G. Dotted lines represent the phase of peak expression according to CircaCompare. For B-G, n=3 biological replicates, ∼100 fly heads per timepoint. For I-J, n=3-6 biological replicates, ∼100 fly heads each per timepoint.

In this study, we investigated the role of *tim* AS in the integration of photoperiod and temperature by the circadian clock to modulate seasonality. We first describe how photoperiod and temperature shape TIM expression. In particular, we show that *tim-sc* is particularly sensitive to changes in photoperiod as compared with *tim-l*, resulting in a significant phase advance in *tim-sc* daily peak expression. TIM-C and TIM-SC are both expressed at the protein level in winter conditions, but TIM-SC is the predominant isoform with regard to expression level. The phase advance observed at the mRNA level is maintained in TIM-SC protein under cold conditions, despite the generally lower amplitude in TIM-SC daily protein rhythm. Interestingly, we found that TIM-SC is predominantly nuclear, affecting circadian clock function and outputs. By performing functional analyses on transgenic flies expressing either *tim-l* or *tim-sc*, we showed that the molecular rearrangement of the core clock incorporating TIM-SC in winter-like conditions allows for a gating mechanism for overwintering. We observed that flies expressing *tim-sc* exhibit low PDF levels, similar to what was previously observed in winter-like conditions^40^, and this coincides with winter-associated daily locomotor profile and reproductive dormancy. Notably, *tim-sc* flies did not completely phenocopy flies maintained in winter conditions with regard to regulation within the molecular clock, suggesting other mechanisms are involved. In summary, we propose a model in which AS-dependent molecular plasticity of clock components is critical to seasonality.

## Results

### Temperature-dependent *tim* isoforms are differentially regulated by photoperiod

*Timeless* mRNA splicing is regulated by temperature^66–69^. At 25 °C, the dominant isoform is the canonical *tim-l*, whereas under cold conditions, two isoforms are produced: *tim-c* and *tim-sc* (Fig. 1A). It is unknown whether photoperiod, as another major seasonal cue, has any effect on the expression of these isoforms at the mRNA level. To answer this question, we determined the levels of *tim-l, tim-c*, and *tim-sc* in heads of wild type flies entrained under summer-like conditions (long photoperiod, 16:8 LD at 25°C) or winter-like conditions (short photoperiod, 8:16 LD at 10°C). We used both Canton-S (CS) flies and *w*^1118^ flies as they carry either the *ls* or *s* allele, respectively^70^. This allowed us to have all possible combinations of *timeless* generated by alternative translation start sites and AS (Fig. 1A). Overall, all *timeless* isoforms are rhythmically expressed in summer and winter, albeit at different expression levels, amplitudes, and with different peak phases of expression (Fig. 1B-G; p value on each graph). In CS, *tim-l* expression is higher in summer as compared to winter (Fig. 1B; MESOR 0.53 vs 0.21, p=2.4e-11) and the phase of its peak expression changes from Zeitgeber time (ZT)16.9 in summer to ZT11.5 in winter (Δ=5.4 hrs, p=9.6e-4). In contrast, the expression of *tim-c* (Fig. 1C) and *tim-sc* (Fig. 1D) increases in winter conditions as compared to summer (MESOR *tim-c*: 0.65 vs 0.50, p=3.13e-16; MESOR *tim-sc*: 0.65 vs 0.01, p=3.8e-18). For *tim-c*, the peak expression in winter is at ZT11.2, while in summer it is at ZT15.9 (Δ=4.7 hrs, p=3.8e-5). For *tim-sc*, in contrast, the peak phase shows a Δ value of around 6.3 hrs, from ZT16.8 in summer to ZT10.5 in winter, but this difference did not reach significance (p=0.72). Similar results were observed in *w*^1118^ flies, with the main difference being that the MESOR in *tim-l* was not significantly different in summer vs winter (MESOR 0.49 vs 0.40, p=0.10), but with a difference in the amplitude of summer vs winter oscillation of *tim-l* (0.39 vs 0.13, p=1.8e-3). While not significant, possibly due to the lower amplitude of its rhythms, the observed Δ in peak *tim-sc* expression was bigger than the one observed in *tim-l* or *tim-c* across seasonal conditions in both CS and *w*^1118^, suggesting a differential regulation of *tim-sc* compared to the other isoforms.

To further investigate if photoperiod alone has any effect on the peak expression of *tim-sc*, newly eclosed *w*^1118^ flies were entrained either in long photoperiod (LP; 16:8 LD), Equinox (12:12 LD), or short photoperiod (SP; 8:16 LD) at constant 25°C (Fig. 1H). Under these conditions, expression of *tim-l* marginally advances around one hour from LP to Equinox, while another hour advance is observed from Equinox to SP but is not statistically significant (Fig. 1I; LP: ZT16.9, Equinox: ZT15.3, SP: ZT14.0, LPvsEqui p.adj=4.5e-2, EquivsSP p.adj=0.12, LPvsSP p.adj=5.2e-3). The peak expression of *tim-sc* shows an advancement from LP to Equinox and, in contrast to *tim-l, tim-sc* shows a sharp advancement from Equinox to SP (LP: ZT16.6, Equinox: ZT15.1, SP: ZT12.5, LPvsEqui p.adj=3.1e-2, EquivsSP p.adj=2.1e-5, LPvsSP p.adj=2.6e-7). This suggests a differential regulation of *tim* isoforms by photoperiod. While the winter-expressed isoform *tim-sc* is extremely sensitive to photoperiod, *tim-l* is largely insensitive to this phenological cue.

To determine if this photoperiodic regulation of *tim* isoforms translates to changes at the protein level, we detected the temperature-dependent isoforms by western blot under simulated summer (16:8 LD at 25°C) vs winter (8:16 LD at 10°C) conditions. Our original antibody against TIM (Rat polyclonal anti-TIM, R5839, RRID: AB_2782953) was generated using the C-terminal segment of the canonical long isoform, which is missing in TIM-C and TIM-SC isoforms (Fig. 1A and Supplementary Fig. 1A-B). To overcome this issue, we developed a new antibody that recognizes the shared N-terminus of the TIM protein (Supplementary Fig. 1A-B). We showed that this new antibody (RRID: AB_3713152) recognizes TIM-L and TIM-SC in fly head lysates (Supplementary Fig. 1C-D). This conclusion was supported by the absence of signal in *tim^0^* flies and staining of FLAG-tagged TIM-SC with α-TIM and α-FLAG (Supplementary Fig. 1E). Additionally, an extra band slightly smaller than TIM-L is observed at 10°C. The size difference and the cold inducibility are consistent with the notion that this is the TIM-C isoform (Supplementary Fig. 1C-D).

Using this new reagent, we detected TIM isoforms in head lysates from CS and *w*^1118^ flies entrained in LP at 25°C or SP at 10°C (Fig. 2A-B). As expected, TIM-L is the predominant isoform in summer, both in CS and *w*^1118^. In winter conditions, TIM-L is replaced by TIM-C and TIM-SC. In CS, TIM-L peaks at around ZT20 (RAIN p=3.75e-15), while in *w*^1118^ its peak expression occurs at ZT24 (RAIN p=3.86e-10) in simulated summer. The roughly 4-hour delay between mRNA and protein peak for TIM-L is expected, given previously reported post-translational regulation of TIM (reviewed in ^73^). Similarly, there was an approximately 4 to 5-hour delay between mRNA and protein peak for TIM-C. Whereas *tim-c* mRNA peaks ∼ZT11.2, TIM-C protein peaks in the middle of the night at ZT16 in both CS and *w*^1118^ (RAIN CS p=3.16e-6; *w*^1118^ p=1.01e-6). This suggests similar post-translational mechanisms may be at work to regulate TIM-C daily abundance. Finally, TIM-SC is not detectable in simulated summer but very abundant in simulated winter. Although TIM-SC was deemed to exhibit daily rhythmicity in both CS and *w*^1118^ (RAIN p=6.94e-4 and p=2.1e-2, respectively), the amplitudes of these rhythms were much weaker than observed for TIM-L. Nonetheless, we observed a clear ∼8-hour phase difference in the expression of TIM-L in summer and TIM-SC in winter in both CS and *w*^111^ flies. Overall, these data suggest that the expression levels of the different *tim* isoforms are regulated not only by temperature but also differentially regulated by photoperiod. In particular, the peak mRNA expression of *tim-sc* exhibits a significant phase advance, which together with post-translational regulation results in an ∼8-hour phase advance at the protein level.

**Figure 2.**
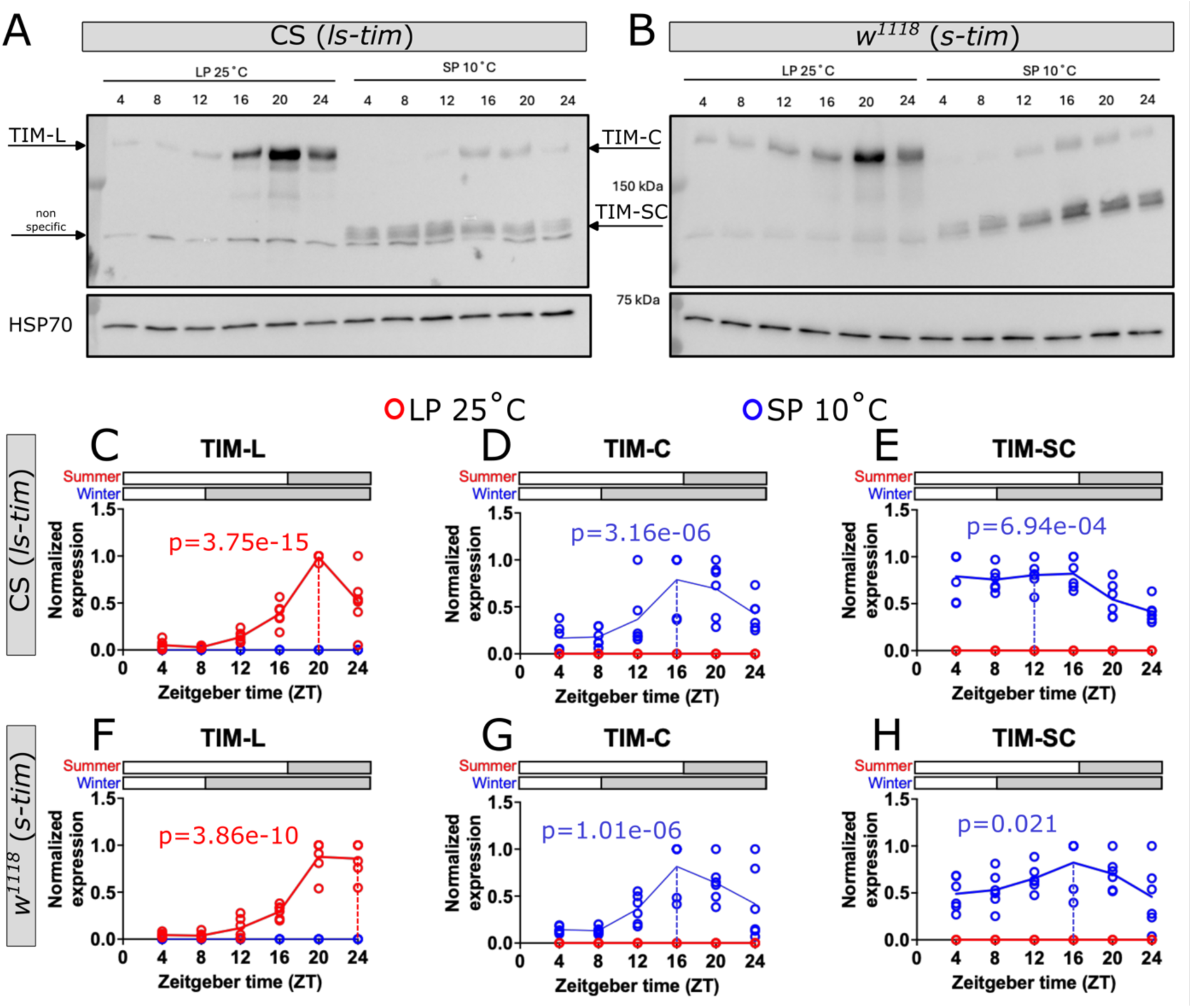
TIMELESS expression differs across different seasonal conditions. Representative western blots detecting TIMELESS using an antibody generated from an N-terminal antigen. Fly head lysates from (A) CS flies, bearing the *ls* allele, or (B) *w*^1118^ flies, bearing the *s* allele, were used. Flies were entrained under summer-like, long photoperiod (LP; 16:8 LD) at 25°C conditions or winter-like, short photoperiod (SP; 8:16 LD) at 10°C conditions, and heads were collected at the indicated timepoints. Arrows indicated the different isoforms based on band size. Quantification of TIM-L, TIM-C, and TIM-SC normalized expression against HSP70, from CS (C, D, E, respectively) and *w*^1118^ flies (F, G, H, respectively). Red lines represent expression at LP 25°C and blue lines represent SP 10°C. Statistical p-values for rhythmicity obtained using CircaCompare are depicted in graphs C-H. The dotted line represents the phase of peak expression of the proteins according to CircaCompare. For C-H, n=6 biological replicates, ∼100 fly heads per timepoint. Horizontal bars above the graphs depict light (white) and dark (gray) periods in summer vs winter conditions.

### TIMELESS is predominantly nuclear in winter conditions

We showed that TIM-SC is the main isoform expressed in whole heads under winter-like conditions. However, whether there are seasonal differences at the cellular level within the circadian clock neuronal network (CCNN) is unknown. In particular, TIM-SC is a truncated protein that is missing the well-characterized cytoplasmic localization domain (CLD) in canonical TIM-L, suggesting that the regulation of its subcellular localization could be compromised. Thus, we sought to determine the expression of TIM across the CCNN under different seasonal conditions, with the understanding that TIM-L is dominantly expressed in simulated summer while TIM-SC is preferentially expressed in simulated winter. Taking advantage of our newly developed antibody, we detected TIM in brains from CS and *w*^1118^ flies entrained under summer conditions (LP 25°C) or winter conditions (SP 10°C). Our antibody readily detects all the major clock clusters, including the Dorsal Neurons 1 (DN1), the Dorsal Neurons (DN3), the Dorsal Lateral Neurons (LNds), and the Ventral Lateral Neurons (LNvs; Supplementary Fig. 2A). Relatively low levels of TIM are observed at ZT3 in summer or winter conditions (Fig. 3A and 3B, respectively). In contrast, higher expression of TIM is observed at ZT15 in both seasonal conditions (Fig. 3C-D). Interestingly, simulated winter appeared to alter the subcellular localization of TIM. In summer, TIM is predominantly cytosolic at ZT15 (Fig. 3C; inserts), while in winter, TIM is almost exclusively nuclear (Fig. 3D; inserts).

**Figure 3.**
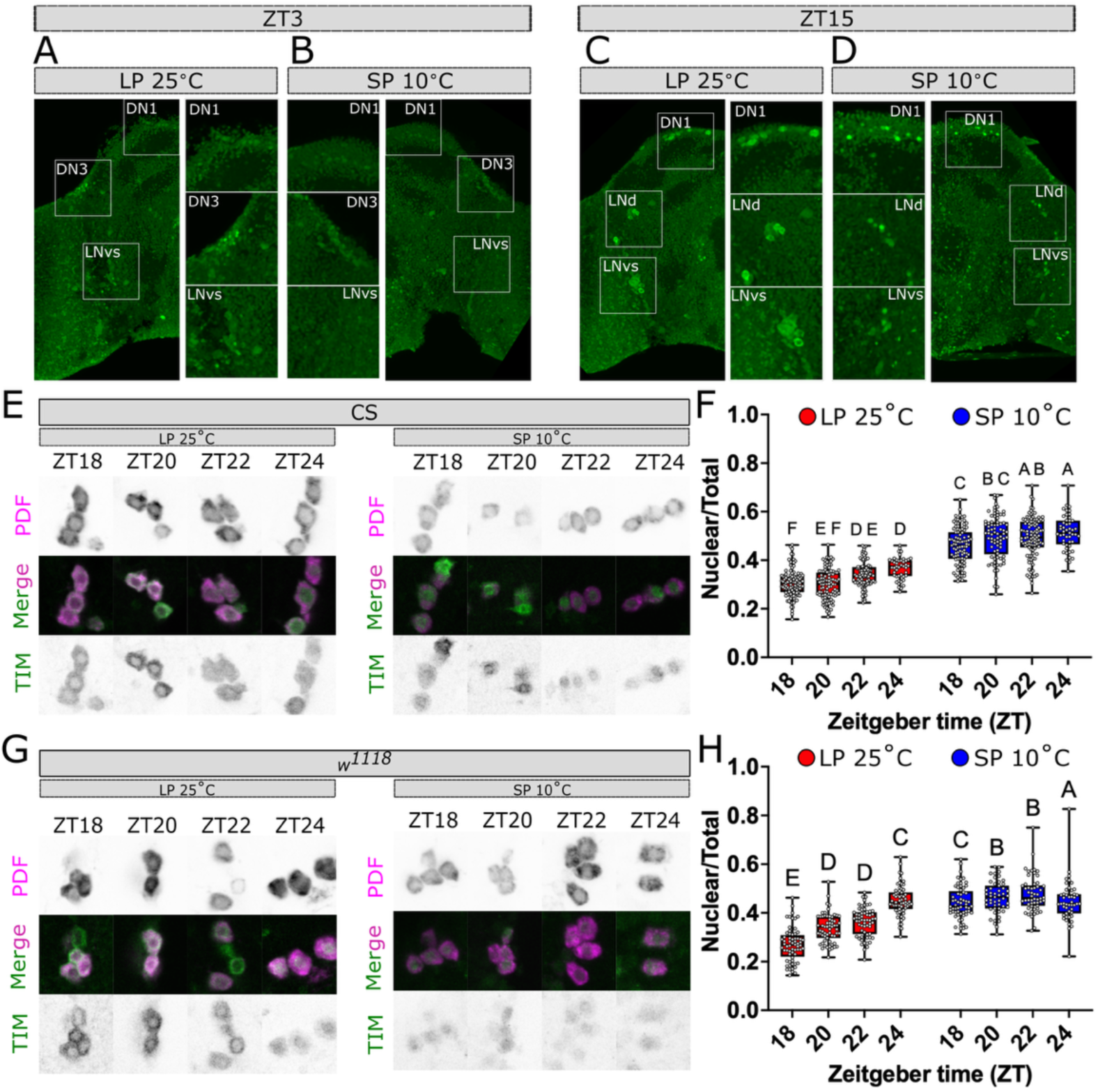
TIMELESS is predominantly nuclear in winter-like conditions. Representative images of TIM expression in brains collected at zeitgeber time 3 (ZT3; A-B) or zeitgeber time 15 (ZT15; C-D) from flies entrained in summer-like, long photoperiod (LP; 16:8 LD) at 25°C conditions or winter-like, short photoperiod (SP; 8:16 LD) at 10°C conditions. The inserts show the main clock neuronal clusters, DN: Dorsal Neurons, LNvs: Ventral Lateral Neurons, LNds: Dorsal Lateral Neurons. Representative images and quantification of co-immunofluorescence against Pigment Dispersing Factors (PDF; magenta) and TIM (green). Samples were collected at ZT18, 20, 22, and 24 from (E-F) CS flies and *w*^1118^ (G-H) entrained in LP 25°C (red boxes) or SP 10°C (blue boxes). Letters represent the significant differences between conditions (different letter: p<0.05, same letter: p>0.05). Two-way ANOVA with Holm-Šídák’s multiple comparisons test, n=45-107 cells from > 8 brains per timepoint, per condition.

To quantitatively determine if the subcellular localization of TIM in summer vs winter conditions, we focused on the LNvs cluster. We used immunodetection of the Pigment Dispersing Factor (PDF) to identify the PDF+ LNvs and to delineate the cytosol and nucleus in brains from CS flies entrained in summer or winter conditions (Fig. 3E). PDF has been used as a cytosol marker in published reports^74,75^. Overall, TIM is more nuclear in winter as compared to summer (Fig. 3F; F(1, 680)=935.3, p<0.0001). Under summer conditions, TIM is mostly cytosolic at ZT18 and progressively becomes more nuclear by ZT24 (0.305 vs 0.367, respectively, p<0.0001). In winter conditions, TIM is already more nuclear at ZT18 as compared to ZT24 in summer (0.459 vs 0.367, p<0.0001). Even with this, TIM nuclear localization increases at later times in winter (ZT18: 0.459 vs ZT24: 0.514, p<0.0001). To explore whether TIM-SC nuclear localization is sensitive to *tim* N-terminal alleles, we conducted the same experiments in *w*^1118^ flies expressing only *s-tim.* Similar results were found in brains from these flies (Fig. 3G). TIM nuclear localization increases with time in both summer and winter (Fig. 3H; F(3, 459)=38.21,p<0.0001) and TIM predominantly nuclear localization is observed in winter as compared to summer (F(1, 459)=229.0, p<0.0001). These data suggest that not only is TIM isoform expression changing across seasons, but so is its subcellular localization.

### TIM-SC expression regulates seasonal locomotor activity

We showed that the expression of TIM isoforms as well as their subcellular localization change across seasons in response to temperature and photoperiod. However, whether these dynamics are important for seasonal adaptations is unknown. Thus, we sought to determine the role of the main TIM isoforms in summer and winter, TIM-L and TIM-SC, respectively, in seasonal adaptations. To do this, we generated two transgenic fly lines expressing either TIM-L or TIM-SC cDNA under ∼4.7 kb of the *tim* promoter (Fig. 4A). As expected, these flies, when used to rescue *tim^0^* mutants, express only TIM-L or TIM-SC at 12:12 LD at 25°C (Fig. 4B). To describe the effect of the selective expression of these isoforms on circadian locomotor activity, flies were kept in 12:12 LD at 25°C for 4 days and then released into constant darkness (DD) for 7 days to assay free-running period. As expected, *yw* control flies show a daily morning and evening peak aligning with the light transitions in LD, which is sustained when flies are placed in DD (Fig. 4C and 4G). Changes in activity at the light transitions are also observed in *tim*^01^ in LD, but no anticipation of these transitions is observed, and rhythms quickly disappear in DD, suggesting a startle response to light instead of rhythms in LD (Fig. 4D and 4H).

**Figure 4.**
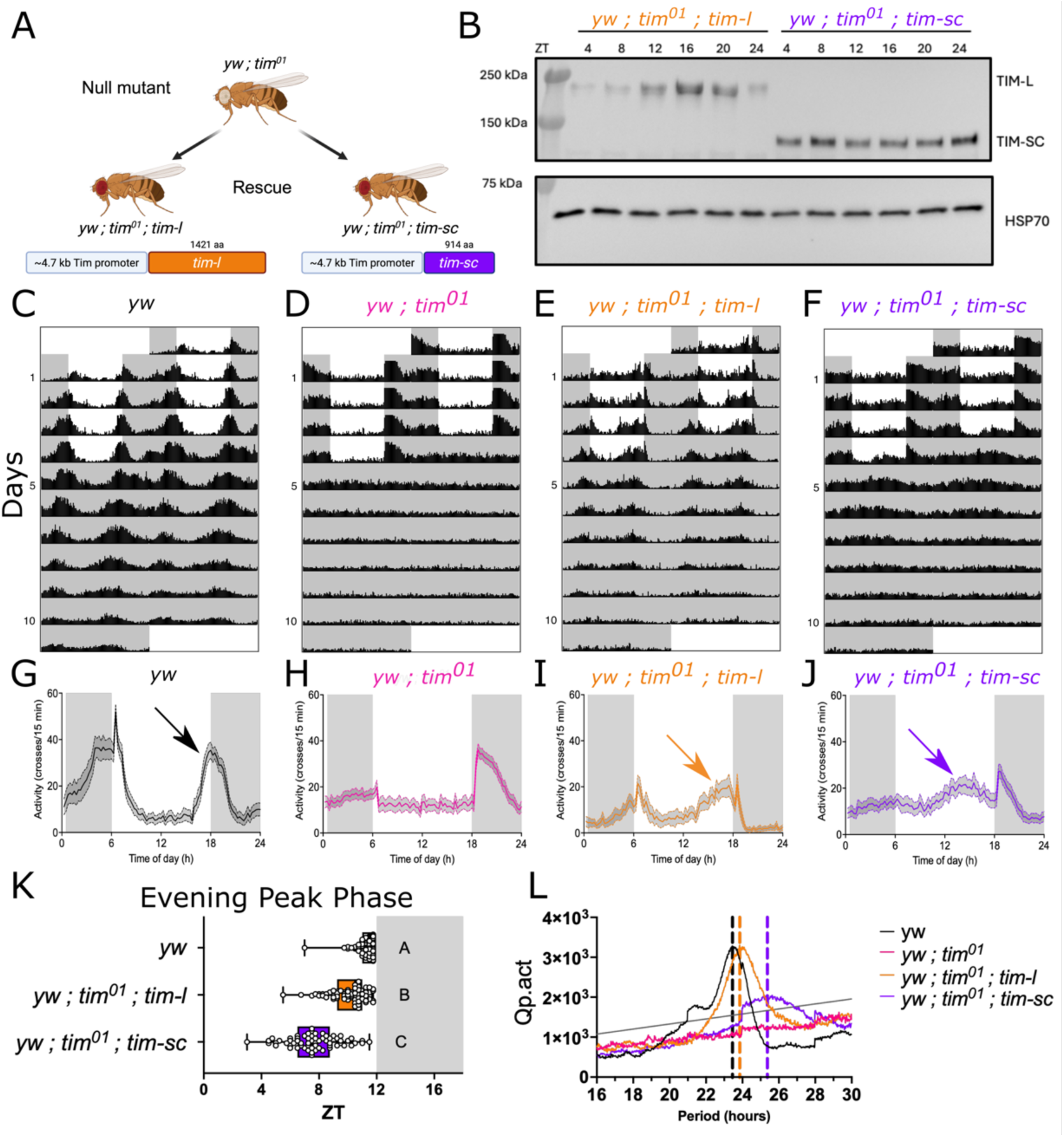
Transgenic-rescue lines expressing specific *timeless* isoforms recapitulate seasonal changes in locomotor activity. (A) Flies expressing only *tim-l* or *tim-sc* were generated by insertion of a transgenic construct bearing the *tim-l* or *tim-sc* coding DNA sequence under a ∼4.7 kb of the *timeless* promoter in *tim*^01^ null-mutant flies. (B) Representative western blot detecting TIMELESS using an antibody generated from an N-terminal antigen in samples from *tim-l* or *tim-sc* rescue flies. Double plot actograms and average activity profiles in LD of *yw* control flies (C, G), *tim^01^* null-mutant flies (D, H), *tim-l* rescue flies (E, I), and *tim-sc* rescue flies (F, J). Grey boxes on the graphs represent the dark period while the light periods are represented as white boxes. The arrow in graphs G-J represent the peak of evening activity. (K) Quantification of the phase of the evening activity peak in *yw* (black box), *tim-l* (orange box), and *tim-sc* (purple box). Letters represent the significant (different letter: p<0.05, same letter: p>0.05) differences between conditions. Kruskal-Wallis test, followed by Dunn’s test. (L) Periodogram for *yw* (black line), *tim* null (pink line)*, tim-l* (orange line), and *tim-sc* (purple line) extracted from the data collected in DD. Dotted lines represent the phase of the peak of Qp.act representing the free-running period for each genotype. The solid line represents the Qp.sig across periods. N=81-87 flies per condition.

One of the main seasonal adaptations observed in *Drosophila* is the plasticity of the evening locomotor activity^40,51,76,77^. During cold days, flies advance their evening activity to potentially match the short days of winter and promote the chances of receiving mid-day heat. While on hot summer days, flies delay their activity to avoid the high heat. Introduction of *tim-l* rescues both daily activity in LD and sustained rhythmic locomotor activity in DD (Fig. 4E and 4I). On the other hand, the introduction of *tim-sc* only recovers the evening peak of activity, while weakly preserving the rhythmicity of this peak on DD (Fig. 4F and 4J, Supplementary Table 1). Quantification of the phase of the evening peak of activity in several flies shows a difference across genotypes (Fig. 4K; H(2)=142, p<0.0001). In particular, there is a 3.6-hour advancement of the peak in *tim-sc* rescue flies as compared to *yw* control flies (7.651 vs 11.28, p<0.0001). This advancement is of similar magnitude to the one observed in *pdf*^01^ flies (Supplementary Fig. S3)^40,78^. A marginal, but significant, 1.17-hour advancement was observed in *tim-l* rescue flies as compared to the *yw* control (10.11 vs 11.28, p<0.0001), while differing in 2.5 hours with the peak of *tim-sc* flies (10.11 vs 7.65, p<0.0001).

Finally, rhythmicity is rescued to the level of *yw* control flies with the introduction of *tim-l* (Fig. 4C-F, Supplementary Table 1), with a period of around 24 hrs (Fig. 4L; black and orange lines, Supplementary Table 1). Rhythmicity is partially rescued by the introduction of *tim-sc*, albeit with a longer free-running period (Fig.4 L; purple line, Supplementary Table 1). All in all, *tim-sc* rescue flies do not behave as *tim*^01^ or *tim-l* flies, suggesting that TIM-SC might have an active function within the clock that differs from the canonical TIM-L isoform.

### *Tim-sc* generates a winter lock to promote successful overwintering

In a previous study, we demonstrated that PDF is a crucial regulator in prompting flies to overwinter^40^. In summer, PDF levels are high, maintaining summer locomotor architecture and promoting reproduction. In winter, a reduction in PDF levels reshapes daily locomotor activity profiles through the modulation of the evening peak and triggers reproductive dormancy. Consistent with the idea that PDF expression can be differentially regulated by *tim* AS, *tim-sc* flies show an advancement in their evening peak of activity similar to *Pdf*-null mutants in 12:12 LD cycles at 25°C (Fig. 4J and Supplementary Fig. 3). To determine if PDF expression is indeed altered in flies expressing *tim-l* vs *tim-sc*, we assayed the levels of PDF in the terminals of the s-LNvs, which are important for seasonal adaptations. As we expected, PDF levels are overall reduced in *tim-sc* flies as compared to *tim-l* flies (Fig. 5A-B; F(1, 103)=30.12, p<0.0001). Additionally, PDF levels are rhythmic in *tim-l* flies while arrhythmic in *tim-sc* flies (Fig. 5B; RAIN p=1.1e-5 and p=0.5, respectively).

**Figure 5.**
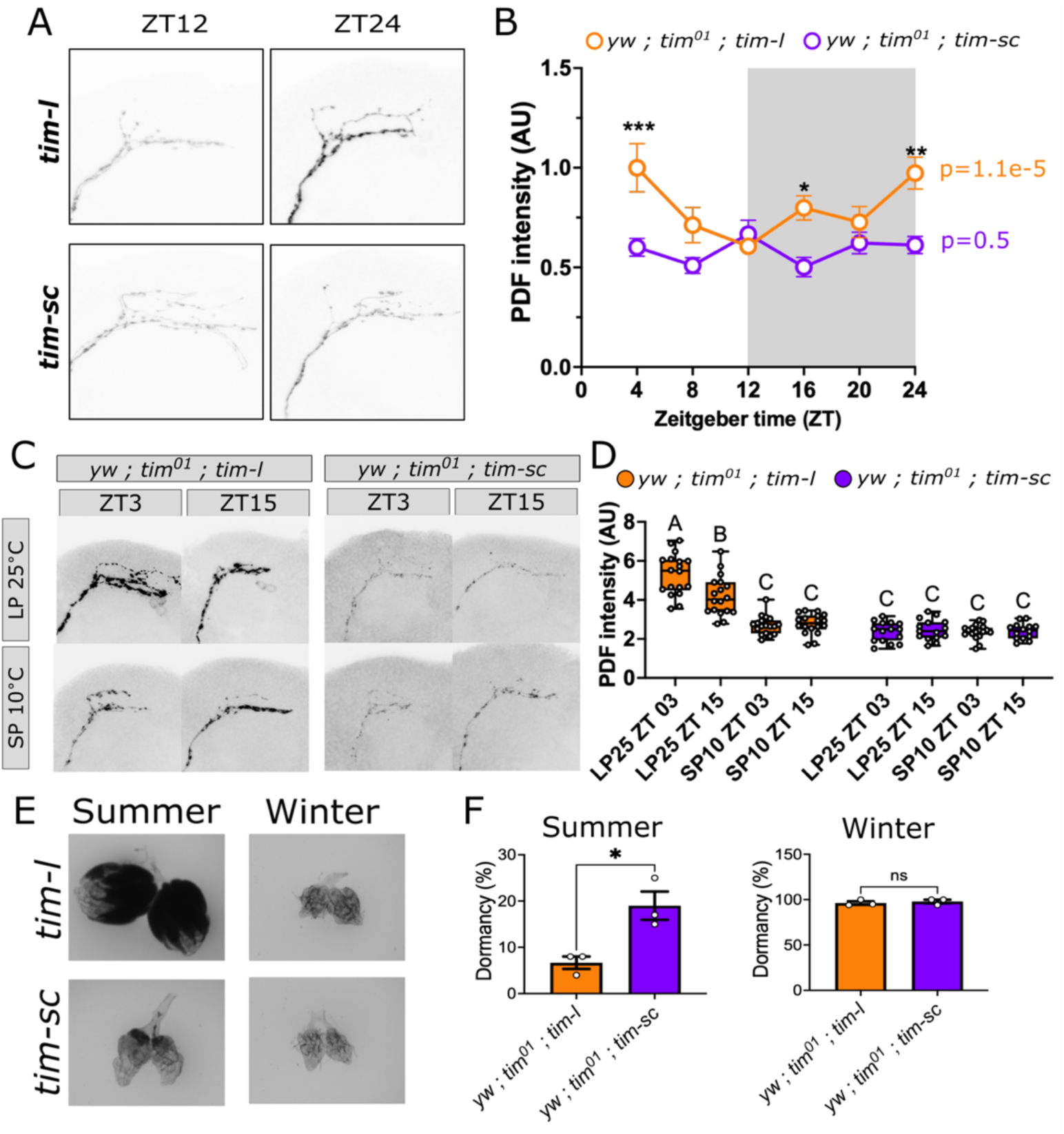
*Timeless* splicing acts as a seasonal lock, to modulate PDF. (A) Representative immunofluorescence of Pigment Dispersing Factor (PDF) in the terminals of the small Ventral Lateral Neurons (s-LNvs) in *tim-l* and *tim-sc* rescue lines. Flies were entrained in 12:12 LD cycles at 25°C for three days and collected at zeitgeber time 12 (ZT12) and ZT24. (B) Quantification of PDF intensity on the s-LNvs in *tim-l* (orange line) and *tim-sc* (purple line) flies. Two-way ANOVA with Šídák’s multiple comparisons test, *p<0.05, **p<0.005, ***p<0.0005, n = 8-10 brains per condition. The p-value for rhythmicity for *tim-l* and *tim-sc* obtained using RAIN is shown in orange and purple letters, respectively. (C) Representative immunofluorescences of PDF in the s-LNvs terminals from flies entrained in summer-like, long photoperiod (LP; 16:8 LD) at 25°C, or winter-like, short photoperiod (SP; 8:16 LD) at 10°C, conditions. Samples were collected at ZT3 and ZT15. (D) Quantification of experiments in panel C. Two-way ANOVA with Holm-Šídák’s multiple comparisons test. Letters represent the significant differences between conditions (different letter: p<0.05, same letter: p>0.05). (E) Representative images of ovaries from *tim-l* and *tim-sc* rescue flies entrained in summer-like conditions (LP 25°C) or winter-like conditions (SP 10°C). (F) Dormancy, as the percentage of flies in a population in ovaries with <1 egg or immature egg, in summer (left panel) or winter (right panel) in *tim-l* (orange bars) and *tim-sc* (purple bars) flies. Unpaired t-test, *p<0.05, n=3 biological replicates with ∼20 flies each.

Considering that PDF levels are low in *tim-sc* flies and high in *tim-l* flies, we hypothesized that *tim* isoform exchange from *tim-l* in summer to *tim-sc* in winter underlies the changes in PDF observed across seasons. To test this, we took *tim-l* and *tim-sc* flies and entrained them under summer (LP 25°C) or winter (SP 10°C) conditions and probed against PDF at ZT3 and ZT15. If *tim* isoform exchange is necessary for PDF-mediated seasonal adaptation, *tim-l* and *tim-sc* flies would be unresponsive to environmental conditions, and PDF would stay high or low, respectively, regardless. However, this is not what we observed. In summer, *tim-l* flies display high levels of PDF that are reduced when placing the flies in winter conditions (Fig. 5C-D; orange boxes; F(3, 126)=28.90, p<0.0001). On the other hand, *tim-sc* flies have low levels of PDF both in summer and winter (Fig. 5C-D; purple boxes). This suggests that *tim* isoform exchange is not required for the reduction of PDF from summer to winter, but instead, the presence of *tim-sc* locks PDF levels in a winter state.

If the winter lock hypothesis is correct, placing *tim-sc* flies even under summer conditions will promote overwintering. To test this, we subjected *tim-l* and *tim-sc* flies to summer or winter conditions and examined ovary development as a readout of reproductive dormancy. Consistent with our hypothesis, *tim-sc* flies have higher levels of dormancy in summer as compared to *tim-l* flies (Fig. 5E-F; left panel; 19% vs 6.7 %; t(4)=3.7, p=0.0208). As expected, both *tim-l* and *tim-sc* enter dormancy in winter-like conditions (Fig. 5E-F; right panel; 96.33% vs 98.00%, t(4)=0.61, p=0.574). In summary, our data show that TIM-SC generates a winter lock, gating PDF in cold, short days to promote a winter program.

### TIM-SC reshapes the molecular clockwork through PER activity

PDF is indirectly regulated by CLK^79,80^. Given the importance of TIM in the molecular clock and its function in regulating CLK activity, the molecular remodeling of the core clock resulting from the expression of TIM-SC in winter conditions may result in the observed reduction of PDF, which underlies the winter lock mechanism. To determine the effect of having TIM-SC instead of TIM-L in the molecular clockwork, we assayed the expression and function of TIM heterodimeric partner PERIOD in *tim-l* and *tim-sc* flies. We first assayed PER levels by immunoblotting in whole head extracts from *tim-l* and *tim-sc* flies (Fig. 6A). As expected, PER levels cycle across the day in *tim-l* flies, with a peak expression at ZT16 (Fig. 6B; RAIN p=5.32e-4). Interestingly, PER expression is significantly reduced in *tim-sc* flies as compared to *tim-l* flies (Fig. 6B; F(1, 60)=47.44, p<0.0001). Additionally, PER daily expression is arrhythmic in *tim-sc* flies (RAIN p=0.91).

**Figure 6.**
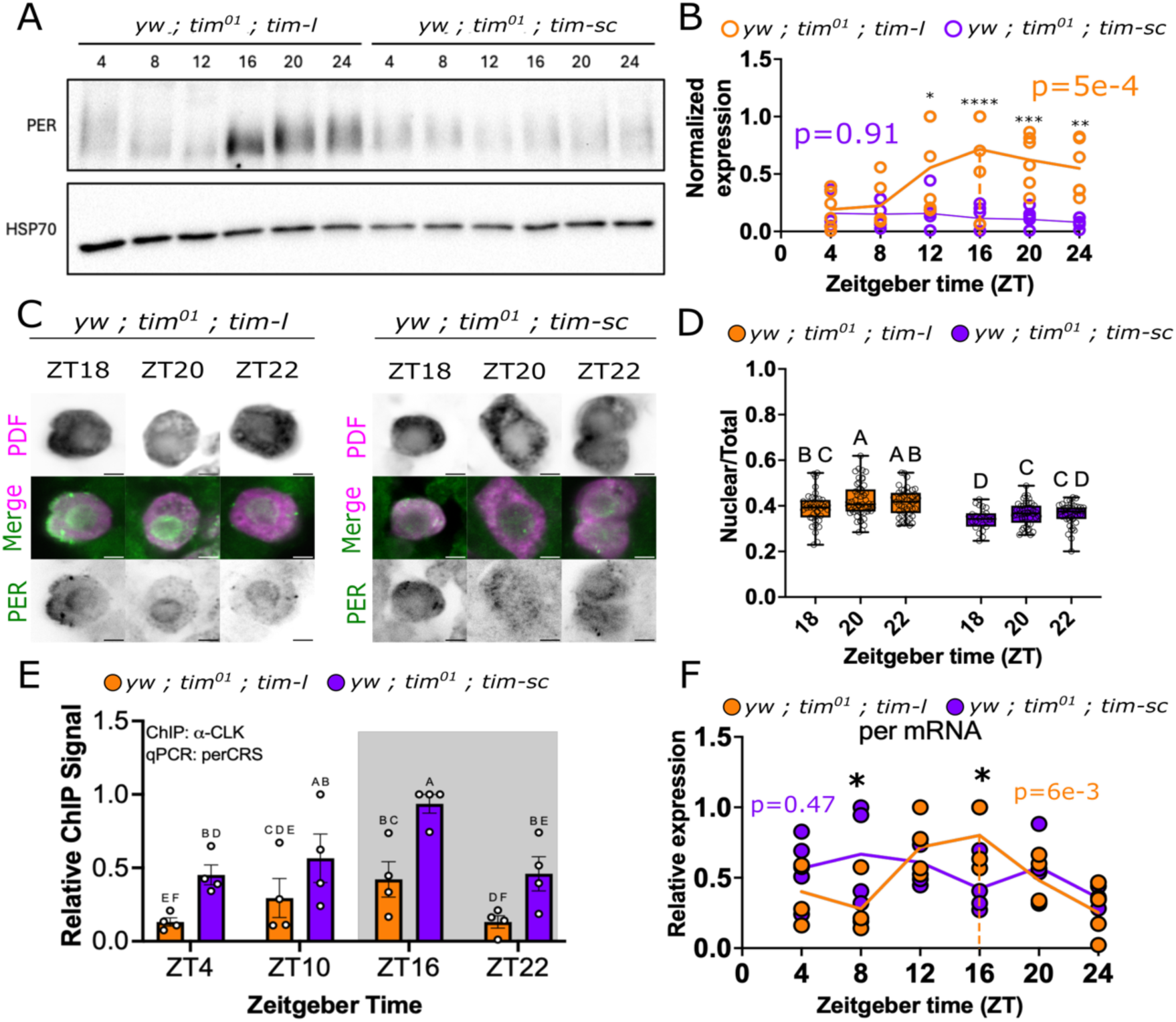
TIM-SC expression affects key core clock features. (A) Representative western blot detecting PERIOD in *tim-l* and *tim-sc* rescue lines entrained in 12:12 LD at 25°C. (B) Quantification of normalized PER expression in *tim-l* (orange line) and *tim-sc* (purple line) fly lines. HSP70 was used for normalization. The dotted line represents the phase of the peak expression, and the p-value represents the rhythmicity statistic according to RAIN. Two-way ANOVA with Šídák’s multiple comparisons test, n=6 biological replicates. *p<0.05, **p<0.005, ***p<0.001, ****p<0.0001. (C) Representative co-immunofluorescence against Pigment Dispersing Factor (PDF; magenta) and PER (green) in *tim-l* and *tim-sc* fly lines. Samples were collected at zeitgeber time (ZT) 18, 20, and 22 in flies entrained in 12:12 LD cycles at 25°C. Two-way ANOVA with Holm-Šídák’s multiple comparisons test, n=23-50 cells from > 8 brains per timepoint, per condition. Letters represent the significant differences between conditions (different letter: p<0.05, same letter: p>0.05). (E) CLOCK occupancy on *per* promoter in heads of *tim-l* (orange bar) and *tim-sc* (purple bar) flies. Two-way ANOVA with Holm-Šídák’s multiple comparisons test, n=4 biological replicates. Letters represent the significant differences between conditions (different letter: p<0.05, same letter: p>0.05). (F) *per* expression on heads from *tim-l* (orange dots) and *tim-sc* (purple dots) rescue lines. Two-way ANOVA with Holm-Šídák’s multiple comparisons test multiple comparisons test, n=4 biological replicates. *p<0.05. The dotted line represents the phase of the peak expression, and the p-value represents the rhythmicity statistic according to RAIN.

A possible hypothesis explaining the reduced PER levels is its altered nuclear localization, resulting in accelerated phosphorylation-dependent degradation. To evaluate subcellular localization of PER, we detected PER by immunofluorescence using a specific antibody (Fig. 6C and Supplementary Fig. 4). As in Fig, 3, we focused on the LNvs by co-staining against PDF. While PER is more nuclear in *tim-l* flies, its distribution in *tim-sc* flies is sparse across the cytosol and the nucleus (Fig. 6C). Quantification of the nuclear fraction of total PER indeed shows a reduction in nuclear localization of the protein in *tim-sc* flies as compared with *tim-l* flies (Fig. 6D; F(1, 234)=45.41, p<0.0001).

Considering that PER localization fails to reach high nuclear levels in *tim-sc* flies, we hypothesized that its downstream function of inhibiting CLOCK-dependent DNA binding and transcription may be affected. To address this, we conducted Chromatin Immunoprecipitation (ChIP), pulling down CLK in *tim-l* and *tim-sc* flies, followed by qPCR against the *perCRS* promoter region (Fig. 6E) to detect CLK-DNA binding. Consistent with our hypothesis, CLK binding to the *per* promoter is significantly elevated in *tim-sc* flies compared to *tim-l* flies (F(1, 27)=25.78, p <0.0001; MESOR p=5.56e-5). Notably, CLK is binding rhythmically to the *per* promoter in both *tim-l* and *tim-sc* flies, regardless of the arrhythmic PER levels in the latter (p=0.02 and p=0.01, respectively). This is in agreement with the rhythmic expression of *tim-c* and *tim-sc* in simulated winter conditions (Fig. 1)

Finally, considering that CLK binding to *per* promoter is higher in *tim-sc* expressing flies, we hypothesized this would result in increased *per* mRNA expression. However, this is not what we observed. Expression of *per* is higher only at ZT8 in *tim-sc* flies (0.667 vs 0.280, p=0.0164), and it is significantly reduced at ZT16, the peak expression of *per* in *tim-l* flies (Fig. 6F; 0.801 vs 0.423, p=0.0193).

Our data suggest that TIM-SC expression affects PER abundance and localization, affecting its role in removing CLK from the DNA. Despite CLK-DNA binding remaining rhythmic in *tim-sc*flies, the expression of *per*mRNA (CLK target) is arrhythmic. This suggests that a partial remodeling of the molecular clockwork may promote seasonal adaptations.

## Discussion

The circadian clock has been functionally linked to seasonal adaptations; however, the exact role of the clock in seasonal timing or to what extent the clock is required to establish, maintain, and exit these adaptations is still unclear. In this study, we demonstrate that the splicing of a core clock component, *tim*, is a crucial factor in maintaining winter adaptations in *Drosophila*. Our data suggest that the presence of TIM-SC in winter, gated by cold temperatures, alters the function of the molecular clock. TIM-SC is primarily present in the nucleus across the day, and this preferential localization promotes PER downregulation, which elevates CLK binding to the DNA. Importantly, this remodeling affects PDF levels, preventing its increase in summer conditions, keeping the animals in reproductive arrest and a constant winter locomotor profile in a process we termed “winter lock” (Fig. 7).

**Figure 7.**
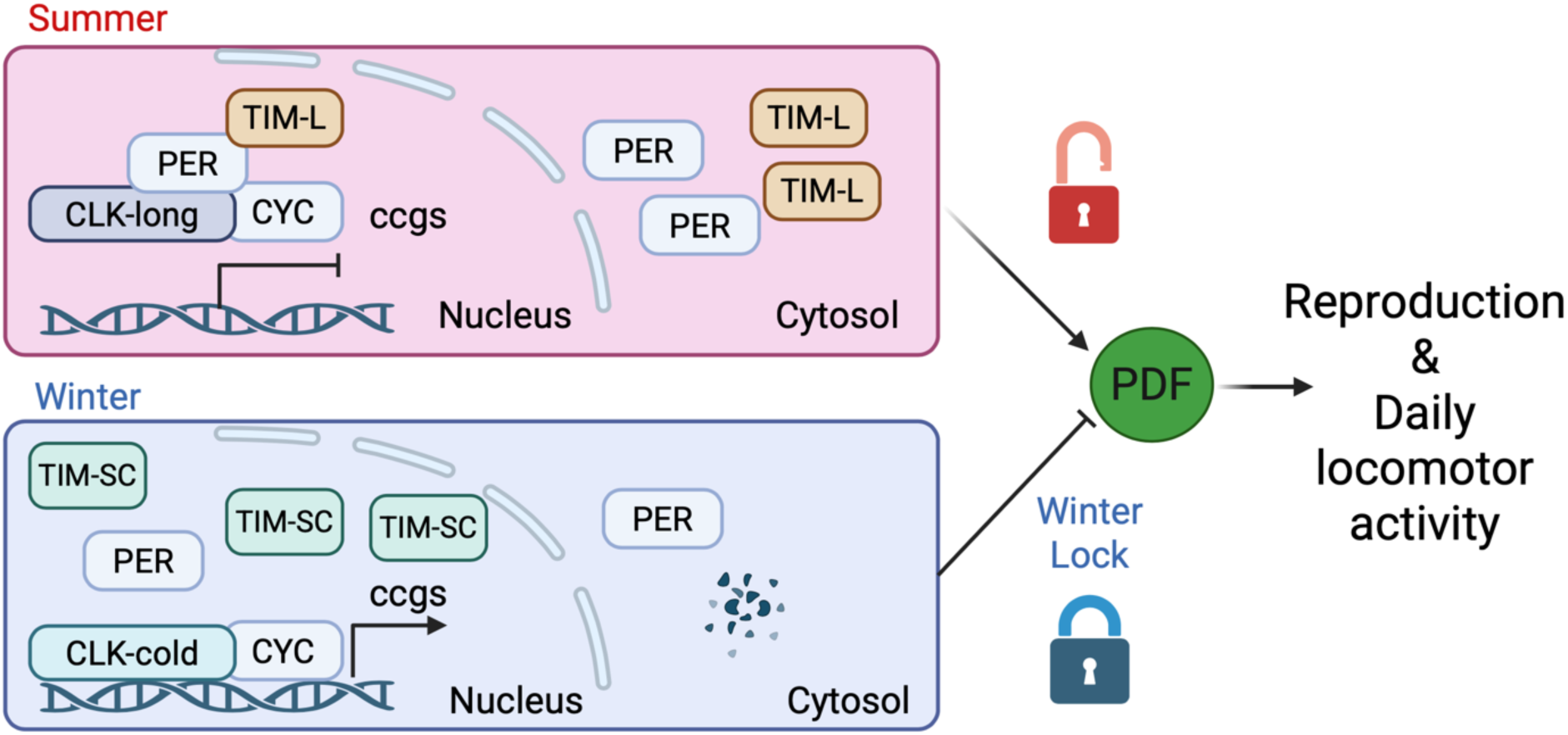
Model depicting the role of *timeless* isoform exchange in seasonal adaptations. During summer, in long photoperiods and warm days, the predominant TIM-L isoform achieves its function of chaperoning PER into the nucleus to inhibit interaction of CLK-long to the DNA. CLK-long is the predominant isoform of CLOCK in warm temperatures^64^. This promotes the normal daily regulation of PDF, permitting reproduction and normal daily locomotor activity. During winter, the predominant TIM-SC isoform, missing the cytoplasmic localization domain, is predominantly localized in the nucleus. Under these conditions, PER levels are downregulated, and the predominant CLK isoform in the cold, CLK-cold, has higher binding to the DNA. The presence of this “cold molecular arrangement” is sufficient to lock the PDF levels in winter-like states, keeping flies in reproductive arrest and locomotion patterns associated with winter. Created with BioRender.com licensed to the lab of J.C.C.

In natural conditions, coinciding environmental signals permit organisms to accurately change their seasonal behavior and physiology. AS offers an important strategy to functionally diversify the genome to respond to these changes. Here we show that the previously described isoform of *tim, tim-sc,* is not only sensitive to temperature but also to photoperiod. Differential regulation of clock gene splicing by different seasonal cues has been observed in other species. In the model plant *Arabidopsis thaliana*, the splicing of core clock genes TOC1 and ELF3 is suppressed by short days, while cold temperature suppresses CCA1 and ELF3 AS, and induces TOC1 splicing^53,81,82^. Interestingly, the AS of *tim* is photosensitive as opposed to thermosensitive in a northern *Drosophila* species, *D. montana*^56^. These highlight the conservation of splicing of clock genes as a general process by which the clock integrates seasonal environmental cues. Whether the splicing of *tim* or other clock components in *D. melanogaster* is affected by other seasonal cues similar to *D. montana,* remains to be explored

While temperature affected the overall expression of *tim-sc*, the phase of its peak expression was regulated by photoperiod (Fig. 1). We have previously identified a similar modulation of PDF and EYES ABSENT expression, at the mRNA and protein levels, by photoperiod and temperature^40^. It is possible, then, that this is a common feature of seasonal cue integration in *D. melanogaster*. Interestingly, the photoperiodic sensitivity we observed in *tim-sc* was not observed in the *tim-l* canonical isoform. Thus, the coincidence of photoperiod and temperature may be important for the regulation of seasonality in *D. melanogaster*. Studies exploring how the coincidence of environmental cues differentially modulates AS are warranted. A recent study showed that seasonal morphs in the butterfly *Bicyclus anynana* display differential splicing that could potentially work with differential expression in modulating seasonal plasticity, further supporting this idea^61^.

At the protein level, we observed a phase advancement of TIM-SC expression. This is likely a combined effect of temperature and photoperiod, and post-translational modifications, accounting for the delay of around 4 hours in the peak mRNA and peak protein levels. Interestingly, we found that the expression of TIM-SC is rhythmic under winter-like conditions. We previously reported that TIM-SC is not rhythmic at 10°C in Equinox^68^. It is possible that the coincidence of low temperatures and short photoperiods enhances rhythmicity of TIM-SC. Our mRNA work supports this notion as *tim-sc* is particularly sensitive to photoperiod.

TIM-L and TIM-C levels are overall low during daytime, while TIM-SC expression, albeit rhythmic, remains high even during the daytime (Fig. 2). This is observed regardless of the N-terminal variations given by the *ls-tim* or *s-tim* alleles, since it is observed in *CS* and *w*^1118^ flies regardless. Thus, the sensitivity of TIM-SC to light may be reduced as compared with the canonical TIM-L. TIM sensitivity to light is mediated by its binding to the photosensitive protein CRYPTOCHROME (CRY)^83–86^. Since the C-terminal CRY-binding domain is missing in the TIM-SC isoform, CRY-TIM interaction and subsequent TIM degradation may be affected. Further experiments would be required to test this directly.

One striking difference we observed was the subcellular localization of TIM in summer and in winter. Under summer conditions, when TIM-L is the predominant isoform, TIM expression is mostly cytosolic during early night and becomes nuclear at later times. In contrast, in winter conditions, when TIM-SC is the more highly expressed isoform, TIM is observed predominantly in the nucleus throughout the night. TIM-SC is missing a CLD, which has been shown to promote TIM retention and accumulation in the cytosol before its interaction with PER and subsequent nuclear translocation^87^. Thus, it is possible that once translated, TIM-SC is immediately translocated into the nucleus, preventing its cytosolic accumulation.

PER abundance and nuclear accumulation rely on TIM function^87,88^. If TIM-SC is actively translocated to the nucleus once it is produced, we would expect to see PER shuttling at the same time. However, this is not what we observed. Flies expressing only *tim-sc* display lower levels of PER, and it is mostly cytosolic. A possibility is that TIM-SC translocates to the nucleus without PER due to a reduced binding affinity as compared with TIM-L. Consistent with this, TIM-SC is missing half of the second PER binding domain, potentially compromising this interaction. A previous study showed that TIM-SC interacts with PER, so other mechanisms could also be at play^67^. Delayed nuclear accumulation of PER and TIM is key for timekeeping, since both PER and TIM accumulate for several hours before translocation^88,89^. This is tightly regulated by kinases, including Casein Kinase 2 (CK2), Casein Kinase 1α (CK1α), DOUBLETIME (DBT)^90–93^, and SHAGGY (SGG)^94–96^. If TIM-SC and PER indeed form a complex, a differential interaction with any of these kinases could contribute to PER’s reduced translocation and/or expression. Additionally, those kinases that translocate with PER and are involved in removing CLK from the DNA, including CK1α, would also be reduced. Further experiments are required to test these hypotheses. Nonetheless, the reduction in nuclear PER is consistent with an overall increase in CLK binding to the DNA, suggesting a reduction in PER-dependent removal of CLK from the DNA. Interestingly, although we observed more CLK binding to the DNA, we did not observe an increase in *per* mRNA as an output. This suggests that on-DNA CLK is still repressed^97^. A possibility is that PER-independent modulation of CLK by CK2 is altered. CK2 extensively phosphorylates TIM and likely interacts and translocates with it, thus modulating CLK activity^93,96^. Further experiments are required to clarify this. Overall, these support the notion that TIM-SC, which is preferentially expressed in winter, remodels the molecular clockwork, producing a winter-clock that differs in function and output from the summer-clock.

Studies in several species, including *D. melanogaster*^98–100^, the bean bug *R. pedestris*, the silk moth *B. mori*^101,102^, the cabbage beetle *C. bowringi*^103^, and the house mosquito *Cx. pipiens*^44,104^, have functionally shown that the expression of core clock genes is required for seasonal adaptations^27,29^. Yet, the extent and direction of effects observed by knocking down or mutating clock genes have been variable. It is possible that the ability of the clock to influence seasonality does not rely on individual genes/isoforms but on the inherent plasticity of the clock to seasonal cues through splicing. Here, we tested this hypothesis by locking the *tim* isoform to the canonical *tim-l* or the winter-predominant *tim-sc*. Flies expressing *tim-sc* have a significant advancement in the evening peak of activity as compared to *tim-l* expressing flies (Fig. 4). This locomotor architecture is similar to the one observed in flies kept in cold conditions^40,105^. Interestingly, *tim-sc* flies do not behave as null-mutants since they have anticipation and, most importantly, they partially rescue circadian rhythmicity consistent with a previous study^67^. This highlights the idea of a remodeling of the clockwork in winter as opposed to a loss of function. We previously showed that the advancement of the evening peak relates to the levels of PDF^40^. In winter, low PDF levels render advancement in the evening peak, similar to *Pdf*-null mutants^78^. Consistently, flies expressing *tim-sc* show low levels of PDF as compared with *tim-l* expressing flies (Fig. 5). This again suggests that *tim-sc* does not result in a *tim* null system, since PDF is constitutively high in *tim*^01^ mutants^106^.

Maintaining or exiting overwintering is key for survival^5^. Here, we showed that locking the plasticity of the clock resulted in impaired seasonal adaptations. While *tim-l* flies are still able to enter into dormancy, regulated by a reduction in PDF, *tim-sc* flies are not able to exit their winter state since no increase in PDF or ovary development was observed when placing the flies in summer conditions. Interestingly, *tim-sc* transcript is not required for the reduction of PDF, but it is required to maintain low PDF levels in winter conditions. Additonally, *tim-sc* flies do not completely phenocopy winter adaptations seen in wild type flies reared in winter conditions^40^. In that sense, other mechanisms might be in place to trigger this PDF reduction. Supporting this notion, other clock components also undergo temperature-dependent AS in *D. melanogaster*, including *per* and *clk*, suggesting a widespread remodel of the molecular clockwork that could also be in play. In *D. melanogaster*, AS of the 3’ terminal intron of *per* is observed under cold temperatures, which contributes to a differential accumulation of *per* affecting seasonal locomotor adaptations ^51,52,57^. We recently showed that *clk* undergoes temperature-dependent splicing, producing *clk-cold*^64^. The product, CLK-COLD, is missing a key residue, Serine 13, that is required for phosphorylation-dependent inhibition of CLK. Mutations of S13 to aspartic acid resulted in flies that did not show a reduction in PDF under cold conditions. Thus, it is likely that *clk* splicing regulates overall PDF levels, while *tim*splicing gates the transition from low to high levels from winter to summer. It is still unknown how this is achieved. It is possible that other CLK outputs, such as HR38 and SR^79^, could be modulating PDF at the transcriptional level, and VRILLE at the post-transcriptional level^80^. Also, whether *clk* splicing isoforms are also affected by photoperiod, and/or how photoperiod and temperature collectively modulate *clk* splicing in the context of seasonal adaptations, remains to be explored. Additionally, the exact interaction among these cold-induced clock components and how they modulate PDF levels in response to temperature changes warrants future research.

In summary, we showed that the AS of a core clock gene, *tim,* is key to locking flies in a winter program. Since maintaining overwintering features is key for survival, the winter lock provided by TIM-SC is perhaps a key adaptation, hence explaining the regulation of *tim* splicing in several insect species^56,67^. Whether similar mechanisms in which splicing of clock components lock the system either in summer or winter are also observed in vertebrates remains to be explored. The core molecular architecture is conserved across species, and splicing of clock components is a common feature for environmental integration in other species^107–109^. Thus, the possibility of splicing or its misregulation mediating the seasonal physiology on health and disease could be explored^110–112^. Our results also support a role of photoperiod in regulating gene expression in *Drosophila*. Future studies exploring the extent of this regulation could provide insights into the regulation of seasonal physiology and behavior by splicing. Finally, this study could shed light on the molecular adaptations to seasons of other insects, including pests and vectors. These features can be exploited to generate insecticides that hijack the splicing of seasonal components to promote mortality by impairing winter adaptations.

## Methods

### Animals

Fly stocks were reared on a standard corn-yeast diet and maintained at 12:12 LD cycles at 25°C. *w*^1118^ flies were obtained from the Bloomington Drosophila Stock Center (BDSC; Stock #3605), and Canton-S (CS) flies were a kind gift from Dr. Nicolás Fuenzalida-Uribe and Dr. Alfredo Guezzi (University of Puerto Rico, from BDSC #64349). *Timeless* rescue flies were generated as indicated below.

### Generation of *timeless (tim)* rescue lines expressing individual *tim* isoforms

To generate *tim-l* expressing flies, pattB-{tim(WT)-3XFLAG-6XHIS} corresponding to the full-length *ls-tim* allele was used as a backbone^93^. PCR mutagenesis was conducted using NEB Q5 Directed Mutagenesis Kit (New England Biolabs) to mutate the second translation start site (s-tim start site; ATG to GGG) to Gly using the following primers: l-tim_mut_F 5’ GAAATCGGTTGGGGACTGGTTAC 3’ and l-tim_mut_R 5’ ACTTTATCAAAGTTCTGATTATTC 3’. The PCR product was used for KLD reaction (KLD enzyme mix, New England Biolabs), and the digested product was used for transformation in DH5α *E. coli*. Transformants were selected, and plasmid DNA was sent for sequencing (Genewiz, Azenta) to confirm the desired mutation. Final pattB{tim-l-3XFLAG-6XHIS} plasmid was sent for injection to BestGene (Chino Hills, CA). To generate the *tim-sc* expressing flies, pattB{tim-l-3XFLAG-6XHIS} was used as a backbone, and the extra C-terminal segment between the end site for *tim-sc* and the end site for *tim-l* sequence was deleted by PCR mutagenesis as described before, using the following primers: TIM-Q5delCterm_R (TIM-SC) 5’ ACCAGGAGCATACCGTTTGG 3’ and 3x FLAG N-term F 5’ GACTACAAAGACCATGACGGT 3’. Both *tim-l* and *tim-sc* transformants were crossed to a *tim*-null mutant background using *tim*^01^ flies^113^ to obtain the rescue flies used in all experiments.

### Total RNA extraction and quantitative RT-PCR

Total RNA extraction and qRT-PCR were performed as previously described^40,64^. Flies were collected on dry ice, and the heads were separated using prechilled sieves. TRI reagent (Sigma-Aldrich, St. Louis, MO) was used to extract total RNA from heads. RNA quantification was carried out on a NanoDrop Spectrophotometer (Thermo Fisher Scientific, Waltham, MA). For reverse transcription, we used the Superscript IV cDNA Synthesis kit (Invitrogen, Waltham, MA) following the manufacturer’s instructions, with 1µg of RNA as starting material. Quantitative PCR was performed on a Bio-Rad CFX96 Real Time System (Bio-Rad, Hercules, CA) using SsoAdvanced Universal SYBR green super mix (Bio-Rad, Hercules, CA) following the following program: initial 95°C for 30 seconds, followed by 40 cycles of 95°C for 5 seconds, and 60°C for 30 seconds. Primers used are as follows: for *tim-l*: 5’ CCCTTATACCCGAGGTGGAT 3’ and 5’ TGATCGAGTTGCAGTGCTTC 3’, for *tim-c*: 5’ GCATCTGTGTACGAAAAGGA 3’ and 5’ ATGTAACCTATGTGCGACTC 3’, for *tim-sc*: 5’ AACACAACCAGGAGCATAC 3’ and 5’ ATGGTCCACAAATGTTAAAA 3’, for *per*: 5’ GACCGAATCCCTGCTCAA 3’ and 5’ GTGTCATTGGCGGACTTC 3’, and for cbp20: 5’ GTCTGATTCGTGTGGACTGG 3’ and 5’ CAACAGTTTGCCATAACCCC 3’, whose expression was used for normalization. Data were analyzed using the ΔΔCt method.

### TIM antibody generation

TIM antibody (RRID: AB_3713152) was generated by Antibodies Inc. (Davis, CA). To generate an antibody against the shared N-terminus of all TIM isoforms, a peptide comprising the region from 292 to 590 amino acids was generated (Supplementary Fig. 1A-B), purified, and injected into rabbits. Serum was processed by affinity purification with TIM peptide, and the purified TIM antibody was validated by immunodetection of TIM in flies, *Drosophila* S2 cells (Supplementary Fig. 1C-E), and in brains for immunofluorescence (Supplementary Fig. 1F-G).

### Protein extraction and Western blot

Protein extraction was carried out as previously described^40,64^. Briefly, the flies were collected and heads separated on dry ice using chilled sieves, which were then used for total protein extraction. Around 150 µL of extraction buffer (0.1% Glycerol, 20mM Hepes pH 7.5, 50mM KCl, 2mM EDTA, 1% Triton X-100, 0.4% NP 40, 1mM DTT, 10µg/mL Aprotinin, 5µg/mL Leupeptin, 1µg/mL Pepstatin A and 0.5mM PMSF) was added for 50µL of heads for each sample, ground using a motorized pestle, and then centrifuged. The supernatant containing the proteins was collected, quantified, aliquoted in SDS loading buffer, and stored at -20°C until electrophoresis. Electrophoresis was carried out in 8% polyacrylamide SDS PAGE gels for TIM and 10% for HSP70, and then proteins were transferred to nitrocellulose membranes (0.45µm, Bio-Rad, Hercules, CA) on a Trans-Blot Semi-dry transfer system (Bio-Rad, Hercules, CA). Membranes were incubated in blocking solution (5% Blotting Grade Blocker Non Fat Dry Milk, Bio-Rad, Hercules, CA) for 1 hour before antibody incubation overnight at room temperature. Membranes were washed in 0.05% TBST (0.05% Tween 20 in 1X TBS) and incubated with secondary antibody for 2 hours before final TBST washes and imaging. For imaging, membranes were incubated with Clarity ECL reagent (Bio-Rad, Hercules, CA) and imaged using a ChemiDoc MP Imaging system (Bio-Rad, Hercules, CA). The dilution of the antibodies is as follows: rabbit α-TIM antibody (RRID: AB_3713152) at 1:3,000, mouse α-HSP70 (Sigma) at 1:7,000, α-PER (GP5620; RRID: AB_2747405) at 1:3,000, α-rabbit-IgG HRP-conjugated (Cytiva, Marlborough, MA) at 1:2,000, and α-mouse-IgG HRP-conjugated (Cytiva) at 1:7,000.

### Chromatin Immunoprecipitation-qPCR (ChIP-qPCR)

ChIP-qPCR was conducted as described in Cai et al.^64^ Fly heads were collected on dry ice and then ground using a chilled mortar and pestle. Nuclear extraction buffer (10mM Tris-HCl pH 8.0, 0.1mM EGTA pH 8.0, 10mM NaCl, 0.5mM EDTA pH 8.0, 1mM DTT, 0.5% Tergitol NP-10, 0.5mM Spermidine, 0.15mM Spermine) was used to homogenize the tissue with a glass dounce homogenizer. After filtration using a 70µm cell strainer and centrifugation, the nuclear fraction was obtained by sucrose gradient. This fraction was used for crosslinking with 0.3% formaldehyde for 10 min, followed by crosslinking quenching with glycine, and then sonicated to fragment the DNA. Samples were centrifuged at 10,000 rpm for 10 min, and the supernatant was processed for pull-down. Guinea pig α-CLK antibody (GP6139, RRID: AB_2827523)^93^ bound to Dynabeads (Thermo Fisher Scientific, Waltham, MA) was used for CLK pull-down. Samples were incubated with the beads for 2 hours at 4°C in IP buffer (50mM Tris-HCl pH 7.5, 2mM EDTA pH 8.0, 1% Triton X-100, 0.1% DOC, 150mM NaCl, 0.5mM EGTA pH 8.0), and then beads were pelleted and washed in washing buffer (50mM Tris-HCl, 1mM EDTA pH 8.0, 1% Triton X-100, 0.1% DOC, 10µg/ml AcBSA (Promega, Madison, WI), 100mM KCl in 1X PBS, 150mM NaCl, 5mM EGTA pH 8.0, 0.1% SDS). Beads were then eluted with elution buffer (50mM Tris-HCl pH 8.0, 10mM EDTA pH 8.0, 1% SDS, 1mM DTT, 50mM NaCl, 4U/ml Proteinase K (NEB, Ipswich, MA), 50µg/ml RNase A (Thermo Fisher Scientific, Pleasanton, CA) and de-crosslinked overnight at 65°C. QIAquick PCR purification kit (Qiagen, Germantown, MD) was used to purify the DNA, which was quantified and used for qPCR. For *perCRS*, we used 5’ TGCCAGTGCCAGTGCGAGTTCG 3’ and 5’ TGCCTGGTGGGCGGCTGG 3’. For background deduction, the average of amplification of two regions was used: 2R intergenic (CP023338) and X intergenic (FBgn0003638). The primers for 2R were 5’ TCAGCCGGCATCATTAGCAGCCG 3’ and 5’ TCGTGTGCGGGAATCTCTGCCG 3’, and for X 5’ ACTGCGTATTCAGGATACATGCC ‘3 and 5’ TGTCCACTTTAATTGATTGCGTGG 3’.

### Drosophila activity monitoring

Daily locomotor activity was monitored using the Drosophila Activity Monitoring System (DAMS, TriKinetics, Waltham, MA) as described previously^40,114^. Data were analyzed using ShinyR DAMS^115^, and actograms were generated with ActogramJ^116^. The evening peak phase was estimated as the highest point of activity of the evening activity peak, as described in previous publications^40^.

### Reproductive Dormancy

Reproductive dormancy assays were conducted as previously described^35,40^. Newly eclosed female flies were collected between ZT4 and ZT10, and placed in incubators (Tritech Research, Los Angeles, CA) in either simulated summer (16:8 LD at 25°C) or simulated winter (8:16 LD at 10°C) for a week. After this period, flies were collected in 70% ethanol, and ovaries were dissected in 1X PBS. Images of individual ovaries were acquired using an EVOS microscope (Thermo Fisher Scientific), and the percentage of dormancy in a population was quantified as the number of ovaries with no eggs or immature eggs over the total. Three trials were conducted, each consisting of ∼20 flies.

### Whole-brain Immunofluorescence

Immunofluorescence was conducted as previously described^40^. Flies were collected at the indicated timepoints/conditions in 4% Paraformaldehyde in 0.2% PBST (0.2% Triton 100 in 1x PBS) and incubated at room temperature for 40 minutes with agitation. After that, the brains were dissected in 0.2% PBST and kept on ice until all conditions were dissected. Dissected brains were re-fixed with 4% PFA for 20 minutes, then washed four times with 0.2% PBST for 10 minutes each. Brains were then incubated in 5% Normal Goat Serum in 0.2% PBST for 30 minutes, followed by overnight incubation with the primary antibody. The brains were then washed four times in 0.2% PBST at room temperature for 10 minutes each. Brains were incubated for 2 hours with the secondary antibodies diluted to their individual working concentration in 5% Normal Goat Serum. Following four 10-minute washes with 0.2% PBST, brains were incubated in 50% Glycerol at 4°C for 30 min. Samples were mounted using ProLong Diamond Antifade Mountant (Fisher Scientific, Waltham, MA) and left in the dark to cure for at least 24 hours before imaging. The dilution of the primary antibodies was as follows: rabbit α-TIM (RRID: AB_3713152) 1:300, mouse α-PDF 1:200 (Developmental Studies Hybridoma Bank), and rabbit α-PER (Rb s3968-1; RRID: AB_2747406) 1:500. The dilution of the secondary antibodies is as follows: Goat α-Rabbit IgG (H+L) Alexa Fluor™ 488 (Invitrogen, Waltham, MA) 1:500 and Goat α-mouse IgG Alexa Fluor® 647 Conjugate (Cell Signaling, Danvers, MA) 1:1,000. PDF C7 was deposited in the DSHB by Justin Blau (DSHB Hybridoma Product PDF C7).

Images were captured with a Leica SP8 confocal microscope equipped with excitation diodes at 638 nm and OPSL exciting at 488 nm. For the experiments determining PDF levels in the s-LNvs dorsal terminals, images were taken every 1 µm using a 40x oil objective and digital zoom. For images of the whole brain, a 20x objective was used instead. All images were analyzed using Fiji, and PDF levels were assessed as the intensity in the s-LNvs dorsal terminal normalized by the background signal in the same dorsal region. To quantify the nuclear and total amount of PER and TIM, images of the LNvs cell bodies were taken every 1µm, and a cross-sectional plane for each cell body was selected for quantification.

### Quantification and Statistical Analysis

For XY graphs, data were represented as mean ± standard error of the mean (SEM). For all other representations, whiskers and boxes were used, and the result of each biological replicate is represented in the graphs as a single dot. For circadian statistics, we used CircaCompare^117^ or RAIN^118^ as indicated for each figure. CircaCompare was used if both conditions in a graph were rhythmic; otherwise, we used RAIN. The specific test used for each dataset is defined in the figure legends, and the statistics for each test are written in the results section with the corresponding p-values. All datasets were tested for normality using a Shapiro-Wilk normality test, and the results were used to inform the best test to use.

## Acknowledgements

The authors want to thank the members of the Chiu lab and the Hamada lab for their comments and feedback. We thank Dr. Pamela Ronald for access to the confocal microscope Leica SP8 from NIH grant GM122968. SH is a Latin American Fellow in the Biomedical Sciences supported by the Pew Charitable Trusts. Research reported in this publication was supported by the National Institute Of Neurological Disorders And Stroke of the National Institutes of Health under Award Number K99NS133470 and R00NS133470. The content is solely the responsibility of the authors and does not necessarily represent the official views of the National Institutes of Health. Research in the lab of J.C.C. is supported by NIH R01 DK124068. The content is solely the responsibility of the authors and does not necessarily represent the official views of the National Institutes of Health.

## Author contributions

SH (investigation, formal analysis, conceptualization, manuscript writing, and editing), CT (investigation, formal analysis, conceptualization, manuscript editing), RDR (investigation, formal analysis, conceptualization, manuscript editing), LAPH (investigation, formal analysis), AB (investigation), YDC (conceptualization, manuscript editing), KL (investigation), JCC (manuscript editing, conceptualization, supervision).

## Additional Information

## Supplementary information

The manuscript contains 1 supplementary table and 4 supplementary figures.

## References

1. Stevenson, T. J., Prendergast, B. J. & Nelson, R. J. Mammalian Seasonal Rhythms: Behavior and Neuroendocrine Substrates. in Hormones, Brain and Behavior 371–398 (Elsevier, 2017). doi:10.1016/B978-0-12-803592-4.00013-4.

2. Jabbur, M. L., Bratton, B. P. & Johnson, C. H. Bacteria can anticipate the seasons: Photoperiodism in cyanobacteria. Science 385, 1105–1111 (2024).

3. J, C., K, O. & T, Y. Light and Hormones in Seasonal Regulation of Reproduction and Mood. Endocrinology 161, (2020).

4. Song, Y. H., Ito, S. & Imaizumi, T. Flowering time regulation: photoperiod- and temperature-sensing in leaves. Trends Plant Sci 18, 575–583 (2013).

5. Denlinger, D. L. Insect Diapause. (Cambridge University Press, 2022). doi:10.1017/9781108609364.

6. Geiser, F. Hibernation. Curr Biol 23, R188–193 (2013).

7. Alerstam, T. & Bäckman, J. Ecology of animal migration. Current Biology 28, R968–R972 (2018).

8. Reppert, S. M., Guerra, P. A. & Merlin, C. Neurobiology of Monarch Butterfly Migration. Annu Rev Entomol 61, 25–42 (2016).

9. Helm, B. & Liedvogel, M. Avian migration clocks in a changing world. J Comp Physiol A 210, 691–716 (2024).

10. Bradshaw, W. E. & Holzapfel, C. M. Evolution of Animal Photoperiodism. Annual Review of Ecology, Evolution, and Systematics 38, 1–25 (2007).

11. Bünning, E. Die endogene Tagesrhythmik als Grundlage der photoperiodischen Reaktion. Berichte der Deutschen Botanischen Gessellschaft (1936).

12. Bünning, E. Circadian Rhythms and the Time Measurement in Photoperiodism. Cold Spring Harbor Symposia on Quantitative Biology 25, 249–256 (1960).

13. Shim, J. S. & Imaizumi, T. Circadian Clock and Photoperiodic Response in Arabidopsis: From Seasonal Flowering to Redox Homeostasis. Biochemistry 54, 157–170 (2015).

14. Hazlerigg, D. G. & Wagner, G. C. Seasonal photoperiodism in vertebrates: from coincidence to amplitude. Trends in Endocrinology and Metabolism 17, 83–91 (2006).

15. Denlinger, D. L., Hahn, D. A., Merlin, C., Holzapfel, C. M. & Bradshaw, W. E. Keeping time without a spine: What can the insect clock teach us about seasonal adaptation? Philosophical Transactions of the Royal Society B: Biological Sciences 372, (2017).

16. Wood, S. H. The Pars Tuberalis and Seasonal Timing. in Neuroendocrine Clocks and Calendars (eds Ebling, F. J. P. & Piggins, H. D.) 33–54 (Springer International Publishing, Cham, 2020). doi:10.1007/978-3-030-55643-3_2.

17. Dardente, H. Circannual Biology: The Double Life of the Seasonal Thyrotroph. Curr Biol 25, R988–991 (2015).

18. Dardente, H. et al. A Molecular Switch for Photoperiod Responsiveness in Mammals. Current Biology 20, 2193–2198 (2010).

19. Dardente, H., Wood, S., Ebling, F. & Sáenz de Miera, C. An integrative view of mammalian seasonal neuroendocrinology. J Neuroendocrinol 31, e12729 (2019).

20. Hazlerigg, D., Lomet, D., Lincoln, G. & Dardente, H. Neuroendocrine correlates of the critical day length response in the Soay sheep. Journal of Neuroendocrinology 30, e12631 (2018).

21. Appenroth, D., Wagner, G. C., Hazlerigg, D. G. & West, A. C. Evidence for circadian-based photoperiodic timekeeping in Svalbard ptarmigan, the northernmost resident bird. Current Biology 31, 2720–2727.e5 (2021).

22. Dupré, S. M. Encoding and Decoding Photoperiod in the Mammalian Pars Tuberalis. NEN G4, 101–112 (2011).

23. Wood, S. & Loudon, A. The pars tuberalis: The site of the circannual clock in mammals? General and Comparative Endocrinology 258, 222–235 (2018).

24. Nakao, N. et al. Thyrotrophin in the pars tuberalis triggers photoperiodic response. Nature 452, 317–322 (2008).

25. Saunders, D. S. Insect photoperiodism: effects of temperature on the induction of insect diapause and diverse roles for the circadian system in the photoperiodic response. Entomological Science 17, 25–40 (2014).

26. Saunders, D. S. & Gilbert, L. I. Regulation of ovarian diapause in *Drosophila melanogaster* by photoperiod and moderately low temperature. Journal of Insect Physiology 36, 195–200 (1990).

27. Hidalgo, S. & Chiu, J. C. Integration of photoperiodic and temperature cues by the circadian clock to regulate insect seasonal adaptations. Journal of Comparative Physiology A 10.1007/s00359-023-01667-1 (2023) doi:10.1007/s00359-023-01667-1.

28. Hamanaka, Y., Hasebe, M. & Shiga, S. Neural mechanism of circadian clock-based photoperiodism in insects and snails. J Comp Physiol A 10.1007/s00359-023-01662-6 (2023) doi:10.1007/s00359-023-01662-6.

29. Goto, S. G. Photoperiodic time measurement, photoreception, and circadian clocks in insect photoperiodism. Appl Entomol Zool 57, 193–212 (2022).

30. Ikeno, T., Numata, H., Goto, S. G. & Shiga, S. Involvement of the brain region containing pigment-dispersing factor-immunoreactive neurons in the photoperiodic response of the bean bug, *Riptortus pedestris*. Journal of Experimental Biology 217, 453–462 (2014).

31. Hamanaka, Y., Yasuyama, K., Numata, H. & Shiga, S. Synaptic connections between pigment-dispersing factor-immunoreactive neurons and neurons in the pars lateralis of the blow fly Protophormia terraenovae. Journal of Comparative Neurology 491, 390–399 (2005).

32. Hasebe, M., Kotaki, T. & Shiga, S. Pigment-dispersing factor is involved in photoperiodic control of reproduction in the brown-winged green bug, *Plautia stali*. J Insect Physiol 137, 104359 (2022).

33. Kotwica-Rolinska, J., Damulewicz, M., Chodakova, L., Kristofova, L. & Dolezel, D. Pigment Dispersing Factor Is a Circadian Clock Output and Regulates Photoperiodic Response in the Linden Bug, Pyrrhocoris apterus. Frontiers in Physiology 13, 1–18 (2022).

34. Colizzi, F. S., Martínez-Torres, D. & Helfrich-Förster, C. The circadian and photoperiodic clock of the pea aphid. J Comp Physiol A 10.1007/s00359-023-01660-8 (2023) doi:10.1007/s00359-023-01660-8.

35. Nagy, D. et al. Peptidergic signaling from clock neurons regulates reproductive dormancy in *Drosophila melanogaster*. PLoS Genetics 15, 1–25 (2019).

36. Lin, Y., Stormo, G. D. & Taghert, P. H. The neuropeptide pigment-dispersing factor coordinates pacemaker interactions in the *Drosophila* circadian system. Journal of Neuroscience 24, 7951–7957 (2004).

37. Shafer, O. T. et al. Widespread Receptivity to Neuropeptide PDF throughout the Neuronal Circadian Clock Network of *Drosophila* Revealed by Real-Time Cyclic AMP Imaging. Neuron 58, 223–237 (2008).

38. Taghert, P. H. & Shafer, O. T. Mechanisms of clock output in the *Drosophila* circadian pacemaker system. Journal of Biological Rhythms 21, 445–457 (2006).

39. Lear, B. C., Zhang, L. & Allada, R. The neuropeptide PDF acts directly on evening pacemaker neurons to regulate multiple features of circadian behavior. PLoS Biology 7, (2009).

40. Hidalgo, S., Anguiano, M., Tabuloc, C. A. & Chiu, J. C. Seasonal cues act through the circadian clock and pigment-dispersing factor to control EYES ABSENT and downstream physiological changes. Current Biology 675–687 (2023) doi:10.1016/j.cub.2023.01.006.

41. Manoli, G., Zandawala, M., Yoshii, T. & Helfrich-Förster, C. Characterization of clock-related proteins and neuropeptides in *Drosophila littoralis* and their putative role in diapause. J of Comparative Neurology 531, 1525–1549 (2023).

42. Manoli, G., Lankinen, P., Bertolini, E. & Helfrich-Förster, C. Correlation between circadian and photoperiodic latitudinal clines in Drosophila littoralis. Open Biol 15, 240403 (2025).

43. Shiga, S. & Numata, H. Roles of PER immunoreactive neurons in circadian rhythms and photoperiodism in the blow fly, *Protophormia terraenovae*. Journal of Experimental Biology 212, 867–877 (2009).

44. Meuti, M. E., Stone, M., Ikeno, T. & Denlinger, D. L. Functional circadian clock genes are essential for the overwintering diapause of the Northern house mosquito, *Culex pipiens*. Journal of Experimental Biology 218, 412–422 (2015).

45. Colizzi, F. S., Veenstra, J. A., Rezende, G. L., Helfrich-Förster, C. & Martínez-Torres, D. Pigment-dispersing factor is present in circadian clock neurons of pea aphids and may mediate photoperiodic signalling to insulin-producing cells. Open Biology 13, 230090 (2023).

46. Cox, K. H. & Takahashi, J. S. Circadian clock genes and the transcriptional architecture of the clock mechanism. Journal of Molecular Endocrinology 63, R93–R102 (2019).

47. Hardin, P. E., Hall, J. C. & Rosbash, M. Feedback of the *Drosophila period* gene product on circadian cycling of its messenger RNA levels. Nature 343, 536–540 (1990).

48. Sehgal, A., Price, J., Man, B. & Young, M. Loss of circadian behavioral rhythms and *per* RNA oscillations in the *Drosophila* mutant *timeless*. Science 263, 1603–1606 (1994).

49. Vosshall, L. B., Price, J. L., Sehgal, A., Saez, L. & Young, M. W. Block in Nuclear Localization of period Protein by a Second Clock Mutation, *timeless*. Science 263, 1606–1609 (1994).

50. Allada, R., White, N. E., So, W. V., Hall, J. C. & Rosbash, M. A Mutant *Drosophila* Homolog of Mammalian Clock Disrupts Circadian Rhythms and Transcription of *period* and *timeless*. Cell 93, 791–804 (1998).

51. Majercak, J., Sidote, D., Hardin, P. E. & Edery, I. How a circadian clock adapts to seasonal decreases in temperature and day length. Neuron 24, 219–230 (1999).

52. Chen, W. F., Low, K. H., Lim, C. & Edery, I. Thermosensitive splicing of a clock gene and seasonal adaptation. Cold Spring Harbor Symposia on Quantitative Biology 72, 599–606 (2007).

53. Kwon, Y.-J., Park, M.-J., Kim, S.-G., Baldwin, I. T. & Park, C.-M. Alternative splicing and nonsense-mediated decay of circadian clock genes under environmental stress conditions in Arabidopsis. BMC Plant Biology 14, 136 (2014).

54. Nilsen, T. W. & Graveley, B. R. Expansion of the eukaryotic proteome by alternative splicing. Nature 463, 457–463 (2010).

55. Xue, R. et al. Alternative Splicing in the Regulatory Circuit of Plant Temperature Response. Int J Mol Sci 24, 3878 (2023).

56. Tapanainen, R., Parker, D. J. & Kankare, M. Photosensitive Alternative Splicing of the Circadian Clock Gene *timeless* Is Population Specific in a Cold-Adapted Fly, *Drosophila montana*. G3 Genes|Genomes|Genetics 8, 1291–1297 (2018).

57. Montelli, S., et al. *period* and *timeless* mRNA Splicing Profiles under Natural Conditions in *Drosophila melanogaster*. Journal of Biological Rhythms 30, 217–227 (2015).

58. Fu, R. et al. Dynamic RNA Regulation in the Brain Underlies Physiological Plasticity in a Hibernating Mammal. Front Physiol 11, 624677 (2020).

59. Jabbour, H. N. et al. Alternative splicing of the prolactin receptor gene generates a 1.7 kb RNA transcript that is linked to prolactin function in the red deer testis. J Mol Endocrinol 21, 51–59 (1998).

60. Tseng, E. et al. Long-read isoform sequencing reveals tissue-specific isoform expression between active and hibernating brown bears (Ursus arctos). G3 (Bethesda) 12, jkab422 (2022).

61. Steward, R. A., de Jong, M. A., Oostra, V. & Wheat, C. W. Alternative splicing in seasonal plasticity and the potential for adaptation to environmental change. Nat Commun 13, 755 (2022).

62. Nakao, N. et al. Photoperiod-Specific Expression of Eyes Absent 3 Splice Variant in the Pars Tuberalis of the Japanese Quail. J Poult Sci 58, 64–69 (2021).

63. Nießner, C. et al. Seasonally Changing Cryptochrome 1b Expression in the Retinal Ganglion Cells of a Migrating Passerine Bird. PLoS One 11, e0150377 (2016).

64. Cai, Y. D. et al. Alternative splicing of *Clock* transcript mediates the response of circadian clocks to temperature changes. Proceedings of the National Academy of Sciences of the United States of America 121, e2410680121 (2024).

65. Cai, Y. D. & Chiu, J. C. *Timeless* in animal circadian clocks and beyond. FEBS Journal 10.1111/febs.16253 (2021) doi:10.1111/febs.16253.

66. Shakhmantsir, I., Nayak, S., Grant, G. R. & Sehgal, A. Spliceosome factors target *timeless (Tim)* mRNA to control clock protein accumulation and circadian behavior in *Drosophila*. eLife 7, 1–27 (2018).

67. Martin Anduaga, A., et al. Thermosensitive alternative splicing senses and mediates temperature adaptation in *Drosophila*. eLife 8, 1–31 (2019).

68. Abrieux, A., et al. EYES ABSENT and TIMELESS integrate photoperiodic and temperature cues to regulate seasonal physiology in *Drosophila*. Proc. Natl. Acad. Sci. U.S.A. 117, 15293–15304 (2020).

69. Boothroyd, C. E., Wijnen, H., Naef, F., Saez, L. & Young, M. W. Integration of light and temperature in the regulation of circadian gene expression in *Drosophila*. PLoS Genetics 3, 0492–0507 (2007).

70. Sandrelli, F. et al. A Molecular Basis for Natural Selection at the *timeless* Locus in *Drosophila melanogaster*. Science 316, 1898–1900 (2007).

71. Tauber, E. et al. Natural Selection Favors a Newly Derived *timeless* Allele in *Drosophila melanogaster*. Science 316, 1895–1898 (2007).

72. Zonato, V., Vanin, S., Costa, R., Tauber, E. & Kyriacou, C. P. Inverse European Latitudinal Cline at the *timeless* Locus of *Drosophila melanogaster* Reveals Selection on a Clock Gene: Population Genetics of *ls-tim*. J Biol Rhythms 33, 15–23 (2018).

73. Zheng, X. & Sehgal, A. Speed control: cogs and gears that drive the circadian clock. Trends in Neurosciences 35, 574–585 (2012).

74. Shafer, O. T., Rosbash, M. & Truman, J. W. Sequential nuclear accumulation of the clock proteins period and timeless in the pacemaker neurons of Drosophila melanogaster. J Neurosci 22, 5946–5954 (2002).

75. Ashmore, L. J. et al. Novel insights into the regulation of the timeless protein. J Neurosci 23, 7810–7819 (2003).

76. Rieger, D., Stanewsky, R. & Helfrich-Förster, C. Cryptochrome, compound eyes, Hofbauer-Buchner eyelets, and ocelli play different roles in the entrainment and masking pathway of the locomotor activity rhythm in the fruit fly *Drosophila melanogaster*. Journal of Biological Rhythms 18, 377–391 (2003).

77. Stoleru, D. et al. The *Drosophila* Circadian Network Is a Seasonal Timer. Cell 129, 207– 219 (2007).

78. Renn, S. C. P., Park, J. H., Rosbash, M., Hall, J. C. & Taghert, P. H. A *pdf* neuropeptide gene mutation and ablation of PDF neurons each cause severe abnormalities of behavioral circadian rhythms in *Drosophila*. Cell GG, 791–802 (1999).

79. Mezan, S., Feuz, J. D., Deplancke, B. & Kadener, S. PDF Signaling Is an Integral Part of the *Drosophila* Circadian Molecular Oscillator. Cell Reports 17, 708–719 (2016).

80. Gunawardhana, K. L. & Hardin, P. E. VRILLE Controls PDF Neuropeptide Accumulation and Arborization Rhythms in Small Ventrolateral Neurons to Drive Rhythmic Behavior in *Drosophila*. Current Biology 27, 3442–3453.e4 (2017).

81. Filichkin, S. A. & Mockler, T. C. Unproductive alternative splicing and nonsense mRNAs: a widespread phenomenon among plant circadian clock genes. Biol Direct 7, 20 (2012).

82. Seo, P. J. et al. A self-regulatory circuit of CIRCADIAN CLOCK-ASSOCIATED1 underlies the circadian clock regulation of temperature responses in Arabidopsis. Plant Cell 24, 2427–2442 (2012).

83. Busza, A., Emery-Le, M., Rosbash, M. & Emery, P. Roles of the Two *Drosophila* CRYPTOCHROME Structural Domains in Circadian Photoreception. Science 304, 1503– 1506 (2004).

84. Emery, P., So, W. V., Kaneko, M., Hall, J. C. & Rosbash, M. CRY, a *Drosophila* clock and light-regulated cryptochrome, is a major contributor to circadian rhythm resetting and photosensitivity. Cell 95, 669–679 (1998).

85. Emery, P. et al. Drosophila CRY is a deep brain circadian photoreceptor. Neuron 26, 493–504 (2000).

86. Naidoo, N., Song, W., Hunter-Ensor, M. & Sehgal, A. A role for the proteasome in the light response of the timeless clock protein. Science 285, 1737–1741 (1999).

87. Saez, L. & Young, M. W. Regulation of Nuclear Entry of the Drosophila Clock Proteins Period and Timeless. Neuron 17, 911–920 (1996).

88. Meyer, P., Saez, L. & Young, M. W. PER-TIM Interactions in Living Drosophila Cells: An Interval Timer for the Circadian Clock. Science 311, 226–229 (2006).

89. Saez, L., Derasmo, M., Meyer, P., Stieglitz, J. & Young, M. W. A Key Temporal Delay in the Circadian Cycle of Drosophila Is Mediated by a Nuclear Localization Signal in the Timeless Protein. Genetics 188, 591–600 (2011).

90. Cyran, S. A. et al. The Double-Time Protein Kinase Regulates the Subcellular Localization of the Drosophila Clock Protein Period. J Neurosci 25, 5430–5437 (2005).

91. Muskus, M. J., Preuss, F., Fan, J.-Y., Bjes, E. S. & Price, J. L. Drosophila DBT Lacking Protein Kinase Activity Produces Long-Period and Arrhythmic Circadian Behavioral and Molecular Rhythms. Mol Cell Biol 27, 8049–8064 (2007).

92. Lam, V. H. et al. CK1α collaborates with DOUBLETIME to regulate PERIOD function in the *Drosophila* circadian clock. Journal of Neuroscience 38, 10631–10643 (2018).

93. Cai, Y. D. et al. CK2 Inhibits TIMELESS Nuclear Export and Modulates CLOCK Transcriptional Activity to Regulate Circadian Rhythms. Current Biology 10.1016/j.cub.2020.10.061 (2020) doi:10.1016/j.cub.2020.10.061.

94. Martinek, S., Inonog, S., Manoukian, A. S. & Young, M. W. A role for the segment polarity gene shaggy/GSK-3 in the Drosophila circadian clock. Cell 105, 769–779 (2001).

95. Ko, H. W. et al. A Hierarchical Phosphorylation Cascade That Regulates the Timing of PERIOD Nuclear Entry Reveals Novel Roles for Proline-Directed Kinases and GSK-3β/SGG in Circadian Clocks. J. Neurosci. 30, 12664–12675 (2010).

96. Top, D., Harms, E., Syed, S., Adams, E. L. & Saez, L. GSK-3 and CK2 Kinases Converge on Timeless to Regulate the Master Clock. Cell Rep 16, 357–367 (2016).

97. Menet, J. S., Abruzzi, K. C., Desrochers, J., Rodriguez, J. & Rosbash, M. Dynamic PER repression mechanisms in the Drosophila circadian clock: from on-DNA to off-DNA. Genes Dev 24, 358–367 (2010).

98. Saunders, D. S., Henrich, V. C. & Gilbert, L. I. Induction of diapause in *Drosophila melanogaster*: photoperiodic regulation and the impact of arrhythmic clock mutations on time measurement. Proceedings of the National Academy of Sciences of the United States of America 86, 3748–3752 (1989).

99. Saunders, D. S. & Bertossa, R. C. Deciphering time measurement: The role of circadian ‘clock’ genes and formal experimentation in insect photoperiodism. Journal of Insect Physiology 57, 557–566 (2011).

100. Pegoraro, M., Gesto, J. S., Kyriacou, C. P. & Tauber, E. Role for Circadian Clock Genes in Seasonal Timing: Testing the Bünning Hypothesis. PLoS Genetics 10, (2014).

101. Zheng, S. et al. Functional polymorphism of CYCLE underlies the diapause variation in moths. Science 388, eado2129 (2025).

102. Ikeda, K., Daimon, T., Shiomi, K., Udaka, H. & Numata, H. Involvement of the Clock Gene *Period* in the Photoperiodism of the Silkmoth *Bombyx mori*. Zoological Science 38, 523–530 (2021).

103. Zhu, L. et al. Circadian clock genes link photoperiodic signals to lipid accumulation during diapause preparation in the diapause-destined female cabbage beetles *Colaphellus bowringi*. Insect Biochemistry and Molecular Biology 104, 1–10 (2019).

104. Meuti, M. E. & Denlinger, D. L. Evolutionary links between circadian clocks and photoperiodic diapause in insects. Integrative and Comparative Biology 53, 131–143 (2013).

105. Miyasako, Y., Umezaki, Y. & Tomioka, K. Separate sets of cerebral clock neurons are responsible for light and temperature entrainment of *Drosophila* circadian locomotor rhythms. Journal of Biological Rhythms 22, 115–126 (2007).

106. Park, J. H., Helfrich-Förster, C., Lee, G., Liu, L., Rosbash, M. & Hall, J. C. Differential regulation of circadian pacemaker output by separate clock genes in *Drosophila*. Proceedings of the National Academy of Sciences of the United States of America G7, 3608–3613 (2000).

107. Liu, Y., Garceau, N. Y., Loros, J. J. & Dunlap, J. C. Thermally regulated translational control of FRQ mediates aspects of temperature responses in the neurospora circadian clock. Cell 89, 477–486 (1997).

108. Diernfellner, A. C. R., Schafmeier, T., Merrow, M. W. & Brunner, M. Molecular mechanism of temperature sensing by the circadian clock of Neurospora crassa. Genes Dev 1G, 1968–1973 (2005).

109. Colot, H. V., Loros, J. J. & Dunlap, J. C. Temperature-modulated alternative splicing and promoter use in the Circadian clock gene frequency. Mol Biol Cell 16, 5563–5571 (2005).

110. Partonen, T. & Lönnqvist, J. Seasonal affective disorder. Lancet 352, 1369–1374 (1998).

111. Rybakowski, J. Forty years of seasonal affective disorder. Psychiatr Pol 58, 747–759 (2024).

112. Zhang, R. & Volkow, N. D. Seasonality of brain function: role in psychiatric disorders. Translational Psychiatry 13, 65 (2023).

113. Myers, M. P., Wager-Smith, K., Wesley, C. S., Young, M. W. & Sehgal, A. Positional Cloning and Sequence Analysis of the *Drosophila* Clock Gene, timeless. Science 270, 805–808 (1995).

114. Cai, Y. D., Hidalgo Sotelo, S. I., Jackson, K. C. & Chiu, J. C. Assaying Circadian Locomotor Activity Rhythm in *Drosophila*. in Circadian Clocks (eds Hirota, T., Hatori, M. & Panda, S.) 63–83 (Springer US, New York, NY, 2022). doi:10.1007/978-1-0716-2577-4_3.

115. Cichewicz, K. & Hirsh, J. ShinyR-DAM: a program analyzing *Drosophila* activity, sleep and circadian rhythms. Communications Biology 1, 25 (2018).

116. Schmid, B., Helfrich-Förster, C. & Yoshii, T. A new ImageJ plug-in ‘actogramJ’ for chronobiological analyses. Journal of Biological Rhythms 10.1177/0748730411414264 (2011) doi:10.1177/0748730411414264.

117. Parsons, R., Parsons, R., Garner, N., Oster, H. & Rawashdeh, O. CircaCompare: a method to estimate and statistically support differences in mesor, amplitude and phase, between circadian rhythms. Bioinformatics 36, 1208–1212 (2020).

118. Thaben, P. F. & Westermark, P. O. Detecting Rhythms in Time Series with RAIN. J Biol Rhythms **2G**, 391–400 (2014).

