## Supplemental Table 1 and Supplemental Figures 1 to 4 for "Splicing of a core clock gene regulates seasonal adaptations by a winter gating mechanism"

Supplementary Table 1. Rhythmicity statistics for isoform-specific genetic rescue of *timeless,* related to Figure 4

| Genotype | Rhythmicity (Qp.act/Qp.sig) | % Rhythmic | Period (h) | n |
|---|---|---|---|---|
| *yw* | 1.10 ± 0.014 | 70.1 | 23.41 ± 0.07 | 87 |
| *yw ; tim*$^{01}$ | 0.907 ± 0.005 | 1.2 | - | 84 |
| *yw ; tim*$^{01}$ *; tim-l* | 1.057 ± 0.016 | 56.8 | 23.67 ± 0.25 | 81 |
| *yw ; tim*$^{01}$ *; tim-sc* | 1.002 ± 0.011 | 33.3 | 24.91 ± 0.39 | 84 |

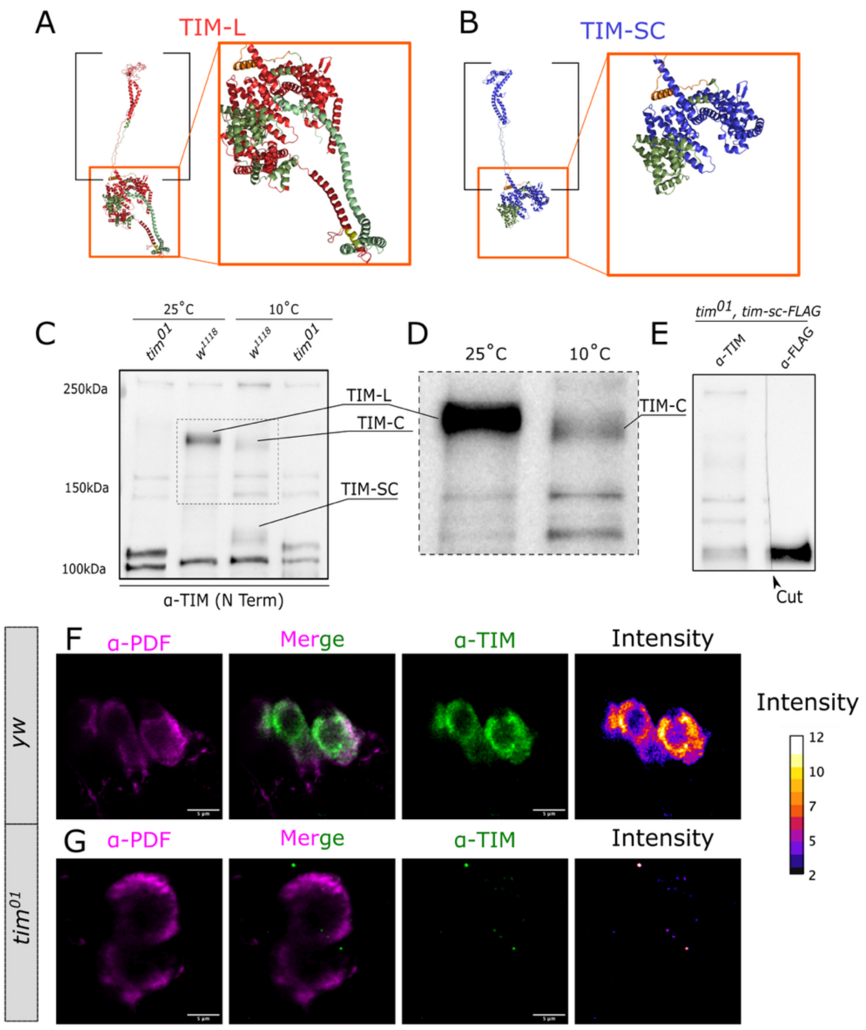


**Figure S1. TIMELESS antibody generation and validation, related to Figure 2 and 3.** Boltz-1 Model of TIM-L (A) and TIM-SC (B). Black brackets enclose the C-terminal segment common to TIM-L and TIM-SC, used to make the antibody. Orange boxes denote the N-terminal domain of the TIM proteins. A zoomed-in version is to the right of panels A and B. In green are the PER binding domains, in orange the Nuclear Localization Signal, and in yellow, in TIM-L, the initial segment of the Cytoplasmic Localization Domain. (C) Representative Western blot of the antibody validation. A band corresponding to TIM-L in $w^{1118}$ is observed at 25˚C that is missing in $tim^{01}$ flies. Bands corresponding to TIM-C and TIM-SC in $w^{1118}$ are observed at 10˚C and are missing in $tim^{01}$ mutants. (D) Contrast-adjusted, zoomed segment of the selected portion of the blot in C. There is a difference in size observed between the two higher molecular weight bands at 25˚C and 10˚C, consistent with the marginal difference in size in TIM-L and TIM-C. (E) Representative western blot of samples from flies expressing TIM-SC-FLAG in a null mutant background. Equal samples were run in parallel, and then the membrane was cut (arrowhead) to probe against TIM and FLAG. All samples were collected at ZT20. (F) Co-immunostaining against Pigment Dispersing Factor (PDF, magenta) and TIM (TIM, green) in *yw* (F) and $tim^{01}$ null-mutant flies (G). Pixel intensity of the TIM channel is observed in the last panels on the right. Scale bar is 5 µm.

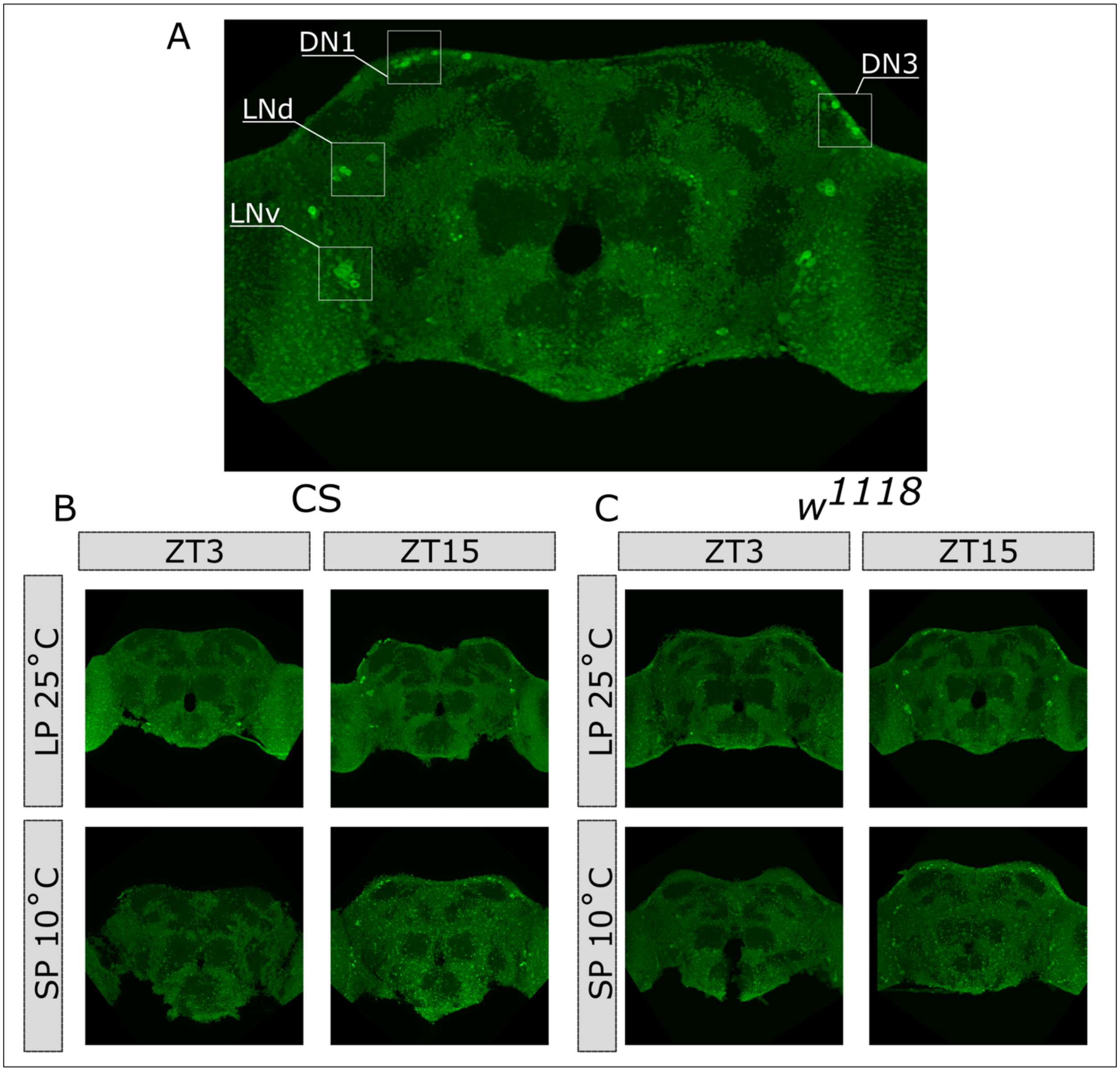


**Figure S2. TIMELESS immunofluorescence, related to Figure 3.** (A) Representative immunofluorescence image detecting TIMELESS in the fly brain using the TIM N-terminal antibody. The circadian clock neuronal cluster Dorsal Neurons 1 (DN1), Lateral Dorsal Neurons (LNds), Dorsal Neurons 3 (3), and Ventral Lateral Neurons (LNvs) are highlighted by the white boxes. Full stack view of brain from Canton-S (CS; B) and $w^{1118}$ (C) in samples collected at ZT3 and ZT15 in flies entrained in long photoperiod (LP; 16:8 LD) at 25˚C or short photoperiod (SP; 8:16 LD) at 10˚C.

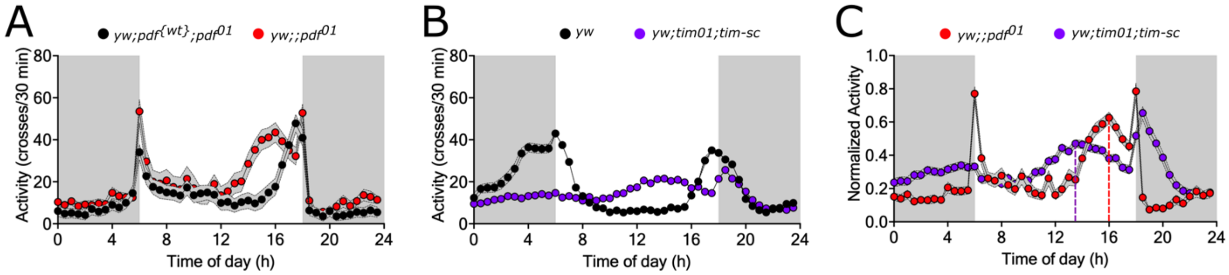


**Figure S3. *pdf*-null mutants and *tim-sc* rescue flies share key changes in locomotor signatures, related to Figure 4.** (A) Average locomotor activity profile of *pdf*-null mutant (*yw* ; *pdf*$^{01}$; red) and *pdf* rescue flies (*yw ; pdf*$^{\{wt\}}$ *; pdf*$^{01}$; black) in 12:12 LD at 25˚C. The phase of the evening peak is advanced in *pdf*$^{01}$ flies. Data reproduced from Hidalgo et al., Current Biology, 2023. (B) Average locomotor activity profile of *tim-sc* rescue flies and yw control flies in 12:12 LD at 25˚C. An advancement of the evening peak is observed in *tim-sc* flies. (C) Normalized activity data from A and B. The phase of the evening peak is observed as a dotted line. N=32 flies per genotype in A and N=96 per genotype in B.

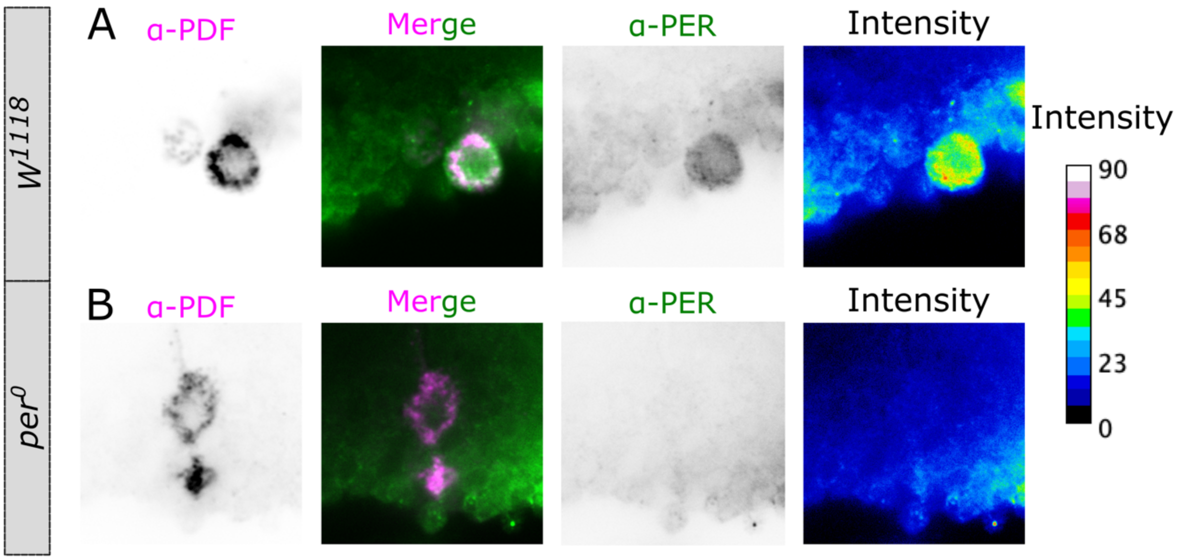


**Figure S4. PER antibody validation for immunofluorescence, related to Figure 5.** Co-immunostaining against Pigment Dispersing Factor (PDF, magenta) and PERIOD (PER, green) in $w^{1118}$ (A) and $per^{0}$ null-mutant flies. Pixel intensity of the PER channel is observed in the last panels on the right.
